# Changes of Cd content in chloroplasts are mirrored by the activity of photosystem I, but not by photosystem II

**DOI:** 10.1101/2023.05.22.541751

**Authors:** Eugene A. Lysenko, Victor V. Kusnetsov

## Abstract

Cd is one of the most toxic heavy metals and widespread pollutant. We searched for a direct Cd action on the photosynthetic electron transport chain using induced chlorophyll fluorescence and P_700_ light absorption. Young barley and maize plants were treated with Cd in toxic (80 μM) and nearly lethal (250 μM) concentrations. The maximal and relative photochemical activities of PSI, its major limitation at the donor side, and partially acceptor-side limitation of PSII changed in agreement with Cd accumulation in the corresponding chloroplasts. Probably, acceptor-side limitation of PSII increased with a direct Cd action under 80 μM that was overcome with an indirect Cd action under 250 μM. These alterations can be explained by Cd/Cu substitution in plastocyanin. The photochemical and non-photochemical quenching by PSII varied diversely that cannot be explained unambiguously by any mechanism. The limitations of PSI (Y(ND), Y(NA)) and PSII (qC) were compared for the first time. They were ranged as follows: Y(NA) < qC < Y(ND). Short segments of qC and Y(ND) dynamics varied proportionally to each other. This implies the existence of an unknown mechanism adjusting limitations at the acceptor side of PSII (qC) and at the donor side of PSI (Y(ND)).

**Highlights:**

- PSI activity changed in agreement with the changes of Cd content in chloroplasts
- The data on PSII activity cannot be clearly explained by Cd action
- PSII acceptor-side limitation qC was governed by opposed direct and indirect Cd actions
- PSI and qC changes can be explained by Cd/Cu substitution in plastocyanin
- Limitations qC of PSII and Y(ND) of PSI changed proportionally for a short time

## 1. Introduction

Cadmium is a highly toxic heavy metal distributed all over the world (Pan et al., 2010; Zou et al., 2021). Sources of Cd accumulation range from natural (e.g. volcanic activity) to industrial and agricultural origin. Mineral fertilizer applications, as well as mining and smelting are considered as the major sources of Cd input into the environment (Bigalke et al., 2017; Kumar et al., 2021). Moreover, Cd mobility is suggested to be very high under natural conditions (Gaillardet et al., 2003; Jigyasu et al., 2020). Many water sources demonstrated relatively high Cd content that is hazardous for both plants and animals (Mahajan et al., 2022).

Plants possess multiple protective mechanisms against Cd toxicity (Seregin and Ivanov, 2001; Lin and Aarts, 2012); however, these mechanisms are mostly energy dependent. The photosynthetic electron-transport chain in thylakoidal membranes is the primary site of energy input into a plant organism. Therefore, protecting chloroplasts against Cd toxicity is of high importance. Most terrestrial plants restrict Cd movement from roots to leaves (Baker, 1981). The small amount of Cd introduces to chloroplasts (Baryla et al., 2001; Pietrini et al., 2003; Lysenko et al., 2015). Inside chloroplasts, Cd is concentrated in thylakoids; in barley thylakoids, the content of Cd is sufficient to inhibit some components of the electron-transport chain (Lysenko et al., 2019).

Plant biologists and photobiologists have been analyzing Cd impact on the photosynthetic electron-transport chain for decades. At present, we have two major problems in this area. First, the data obtained have demonstrated a controversy of Cd impact on the processes around the photosystem II (PSII). The maximal quantum yield (Fv/Fm) of PSII is rather resistant to Cd action. In many studies, Cd treatment did not change Fv/Fm (Burzynski and Zurek, 2007; Shi and Cai, 2008; Lysenko et al., 2015); however, the fall of Fv/Fm was also demonstrated (Dias et al., 2013). The actual quantum yield Փ_PSII_ (synonym Y(II)) and photochemical coefficient qP are more sensitive to Cd action. The Cd treatment decreased both Փ_PSII_ and qP simultaneously (Ci et al., 2010) or only one of these coefficients while the second one remained rather unchanged; the reduced coefficient was Փ_PSII_ (Burzynski and Zurek, 2007) or qP (Lysenko et al., 2020). In a Zn/Cd hyperaccumulator plant, Cd treatment influenced neither Փ_PSII_ nor qP (Tang et al., 2013). Using the new coefficient X(II), an increase in PSII photochemical activity was also demonstrated (Lysenko et al., 2020). Non-photochemical quenching of light energy by PSII was diversely regulated. Cd treatment increased (He et al., 2008), did not change (Burzynski and Zurek, 2007), and decreased (Lysenko et al., 2015; 2020) non-photochemical quenching. We have considered experiments with rather high Cd concentrations: of about 50 μM or similar.

Second, the changes observed by researchers can be caused by either direct or indirect Cd action. Discriminating between direct and indirect mechanisms of Cd action is the fundamental question; we have analyzed this problem in the introduction of previous article (Lysenko et al., 2019). Two more features of this area are rare analysis of the photosystem I (PSI) activity and of Cd accumulation inside chloroplasts.

We have found a model allowing us to distinguish between direct and indirect action of Cd on processes in chloroplasts (Lysenko et al., 2015). In this work, CdSO_4_ was introduced to a mineral media to the final concentrations 80 μM and 250 μM. For both maize and barley plants, the concentration 80 μM was quite inhibitory and far from lethal; the concentration 250 μM was very poisonous and lethal in the prolonged experiment (Klaus et al., 2013; Lysenko et al., 2015). Consequently, elevating Cd concentration from 80 μM to 250 μM should increase numerous indirect effects similarly in both species.

At 80 μM CdSO_4_, the maize plants accumulated a small amount of Cd in chloroplasts (49 ± 5 ng/mg Chl); under 250 μM CdSO_4_, the maize plants acquired more Cd in chloroplasts (92 ± 20 ng/mg Chl) (Lysenko et al., 2015). Therefore, elevating the concentration from 80 μM Cd to 250 μM Cd should increase a direct effect of Cd in the chloroplasts of maize. The barley plants showed an unexpected pattern of Cd accumulation. At 80 μM CdSO_4_, the barley plants accumulated a large amount of Cd in the chloroplasts (171 ± 26 ng/mg Chl). At 250 μM CdSO_4_, the accumulation of Cd rather decreased both in the leaves (significantly) and chloroplasts (126 ± 7 ng/mg Chl) (Lysenko et al., 2015). The latter decrease was insignificant; hence, we assume no further increase of Cd content in the barley chloroplasts at 250 μM CdSO_4_. Therefore, the concentration growth from 80 μM Cd to 250 μM Cd should not increase a direct effect of Cd in the chloroplasts of barley. Thus, we can differentiate Cd actions on the processes in photosynthetic electron-transport chain. At 250 μM CdSO_4_, a similar increase of Cd impact in both species should be considered as an indirect effect. A direct effect may be suggested if the application of 250 μM CdSO_4_ increases Cd impact in maize and does not change Cd effect in barley.

We have revealed an unusual effect in barley plants treated with 80 μM Cd. The photochemical activity of PSII was limited at the acceptor side as indicated by a high level of closed PSII complexes; the photochemical activity of PSI was reduced more than the photochemical activity of PSII (Lysenko et al., 2020). The effect was specific for Cd (Lysenko et al., 2020). These results were obtained with the use of pulse amplitude modulation (PAM) technique, applying the method of rapid light curves (RLC). These changes in the photochemical activities of PSII and PSI could have resulted from either direct or indirect Cd action. To distinguish between direct and indirect Cd action, it is beneficial investigating plants with different Cd accumulation in chloroplasts. The pair of barley and maize represents a favorable experimental model enabling us to differentiate between direct and indirect Cd action on processes in thylakoidal electron-transport chain.

The present study was carried out according to the previous experiment that defined Cd content in the chloroplasts (Lysenko et al., 2015). Barley and maize plants were treated with 80 μM and 250 μM CdSO_4_ for six days; nine-day-old plants were analyzed with Dual-PAM-100. The high level of closed PSII and the inhibition of PSI activity were revealed with the RLC method (Lysenko et al., 2020). In the current study, the photochemical activities of PSII and PSI were investigated with both classic induction curves (IC) and RLC methods. The thorough analysis of PSII and PSI activities and their ratio is presented below. The ratios between limitations of PSII (qC) and PSII (Y(ND), Y(NA)) were calculated for the first time.

## 2. Material and methods

### 2.1. Plant growth conditions

Barley (*Hordeum vulgare* L. cv. Luch) and maize (*Zea mays* L. cv. Luchistaya) seedlings were grown in phytotron chambers at 21°C (barley) or 25-26℃ (maize), 180–220 µmol photons m^-2^ s^-1^ and a photoperiod of 16 h light/8 h dark under continuous aeration on modified Hoagland medium (Lysenko et al., 2019). Preliminary, caryopses were kept for 2-3 days at 4 °C in the dark on a filter paper moistened with 0.25 mM CaCl_2_; imbibed caryopses were germinated in the growth conditions. Two-day-old seedlings were transferred to vessels with Hoagland medium. The next day, CdSO_4_ was added to the hydroponic media to the final concentrations 80 µM or 250 µM; CuSO_4_ was added to the final concentration 80 µM; Fe SO_4_ was added to the final concentration 1.5 mM. The analyses were performed on nine-day-old seedlings. All experiments were repeated at least three times. For more details see (Lysenko et al., 2015; 2019; 2020).

### 2.2. PAM analysis

The chlorophyll (Chl) *a* fluorescence and P_700_ light absorption were simultaneously registered with the Dual-PAM-100 (Walz, Germany). The largest fully developed leaf was used for the measurement: the first leaf of barley and second leaf of maize. The Chl *a* fluorescence was excited at 460 nm (Int. #5, 12 μmol photons m^-2^ s^-1^; the measuring light). P_700_ is the reaction center chlorophyll of PSI; the level of oxidized P_700_ was measured as the difference in light absorption at 830 and 875 nm (Int. #5). The red light 635 nm was used as the actinic light (AL) and for the saturation pulses (SPs).

The plants were adapted in the dark for 30 min, transferred rapidly to the Dual-PAM-100, avoiding bright light, and kept in the dark for another 4 min. In the dark, the measuring light only was used for the determination of minimal (Fo) Chl fluorescence; the first SP (4 mmol photons m^-2^ s^-1^, 500 ms) was applied for the measurement of maximal (Fm) Chl fluorescence; the pre-illumination with far-red light (720 nm, Int. #10) followed with the second SP were employed for the determination of minimal (Po) and maximal (Pm) P_700_ absorption. The measurement of basic parameters in the dark-adapted state was followed with the extra 30 s of darkness.

Next, AL (128 μmol photons m^-2^ s^-1^) was turned on; IC was measured for 7.5 min. The first SP was applied after 0.5 s since AL induction; then, SPs were applied every 40 s. The maximum level of fluorescence in the light (Fm') and maximal P_700_ change in the light (Pm') were measured during SP; the steady state fluorescence (Fs, synonym F) and P_700_ light absorption (P) were measured just before SP. After SP, the illumination was switched to the far-red light (720nm, Int. #10) for 5 s and the minimum level of fluorescence in the light (Fo') was determined. The minimal level of P_700_ light absorption (Po) was measured after cessation of far-red light both in the dark and light regimes (Klughammer and Schreiber, 1994, 2008).

The method IC is the correct and, probably, preferable way of plant adaptation to light conditions before an application of RLC method (Lysenko, 2021). After the completing of IC, the method RLC was started in 6 s (software's switching time). In RLC, each step of AL lasted for 30 s; the set of AL intensities is shown in the figures. The other parameters were the same as in IC.

The parameters of P_700_ light absorption were calculated as follows: Y(I) = (Pm' – P)/(Pm – Po); Y(ND) = (P – Po)/(Pm – Po); Y(NA) = (Pm – Pm')/(Pm – Po); Y(I) + Y(ND) + Y(NA) = 1 (Klughammer and Schreiber, 1994, 2008).

The parameters of Chl fluorescence were calculated using the following equations: Fv = Fm – Fo; Fv' = Fm' – Fo'; *Φ*_PSII_ = (Fm' – Fs)/Fm' (Genty et al., 1989; Kalaji et al., 2014); qP = (Fm' – Fs)/Fv', qN = (Fv – Fv')/Fv (Schreiber et al., 1986; van Kooten and Snel, 1990); X(II) = (Fm' – Fs)/Fv, qC = (Fs – Fo')/Fv (Lysenko et al., 2020); NPQ = (Fm – Fm')/Fm'; (Bilger and Björkman, 1990); qN + X(II) + qC = 1 (Lysenko et al., 2020).`#`

### 2.3. Calculation of ratios between Chl fluorescence and P_700_ absorption

We calculated the ratios between the photochemical activities of both photosystems (Y(I)/X(II)) (Lysenko et al., 2020) and between their limitations (qC/Y(NA) and qC/Y(ND)). In some cases, the values of qC, Y(NA), and Y(ND) were very small. In case when such small value occurred in the denominator of the ratio then it should be removed. The zero values must be discarded because it is forbidden to divide by zero. The negative values have to be discarded because dividing by a negative number generates a nonsense value. The tiny positive values originated extremely high ratios; a single enormous ratio is able distorting a mean of 20-30 normal ratios by several times. For the last case, we have established the criteria for eliminating these data point.

A small signal comparable to a baseline fluctuation can be computed as negative, zero, or extremely small positive. Therefore, the values of denominators (Y(NA) and Y(ND)) were considered. Values qC are frequently low; therefore, qC was used as numerator. Denominator values ≤ 0.025 can be discarded when they are outside of relatively continuous distribution. Denominator values 0.025 < X > 0.030 can be discarded a) if their ratios were 2 times higher than largest point of relatively continuous distribution or b) if their ratios were 1.8 times higher than largest point of relatively continuous distribution, however, removal of this single data point decreased standard error (SE) at least by one third. Denominator values ≥ 0.030 can be discarded a) if their ratios were 3 times higher than largest point of relatively continuous distribution or b) if their ratios were 2.5 times higher than largest point of relatively continuous distribution, however, removal of this single data point decreased SE at least by one half. Some suspicious data points can not be discarded according to these criteria; they can prevent elimination of larger, apparently wrong ratios. One or two “irremovable” data points were considered as outliers; three “irremovable” data points were considered as an edge of relatively continuous distribution. These threshold criteria have been selected empirically. The qC/Y(ND) data points were discarded in the beginning of curves only (40 s in IC, AL 72 and 97 μmol photons m^-2^ s^-1^ in RLC). The qC/Y(NA) data points were discarded from any part of curves; if half or more of data points were removed from a curve then this curve was discarded completely. The portion of data points discarded was relatively small; in a single point, half of data points were removed.

The values Y(I) and X(II) demonstrated stable ratios; seven denominator data points were removed from all the data set. In the beginning of IC (AL 0.5 s) and at the final of RLC (AL 1954 μmol photons m^-2^ s^-1^), the level of X(II) is very low; the values X(II) ≤ 0.005 in barley and X(II) ≤ 0.001 in maize were discarded.

The general map of the data points discarded is represented in Suppl. Tables S1-S3.

### 2.4. Statistics

Control plants and plants treated with 80 μM Cd were studied in three independent experiments. Because of greater variability under nearly lethal conditions 250 μM Cd, plants were studied in four complete independent experiments; for barley plants, IC were measured in two more experiments. The numbers of plants measured with the methods “Darkness+IC / RLC” were as follows: barley 17/17 in control, 17 /17 at 80 μM Cd, and 37/21 at 250 μM Cd; maize 18/18 in control, 18/18 at 80 μM Cd, and 24/19 at 250 μM Cd. Additional treatment with 80 μM Cu and 1.5 mM was performed in six independent experiments; IC were measured in 36 and 37 barley plants correspondingly.

The data were processed using the Excel (Microsoft) software. The significance of differences between mean values was verified using the two-tailed Student's *t*-test.

## 3. Results

### 3.1. Maximal Chl fluorescence and P_700_ absorption in darkness

Parameters measured in the dark-adapted state show maximal activities of the photosystems with no limitations; they therefore reflect the quantitative levels of both photosystems. The barley plants demonstrated a gradual decline of all the parameters of Chl fluorescence: Fm, Fv, Fo, and Fv/Fm (Fig. 1); the decrease was only significant at 250 μM Cd. In contrast, P_700_ light absorption correlated with Cd content in the chloroplasts. At 80 μM Cd, both Pm and Pm/Fv decreased substantially; the larger concentration 250 μM Cd enlarged neither Cd content in the chloroplasts (Lysenko et al., 2015), nor Cd effect on Pm and Pm/Fv (Fig. 1D,F).

**Fig. 1.**
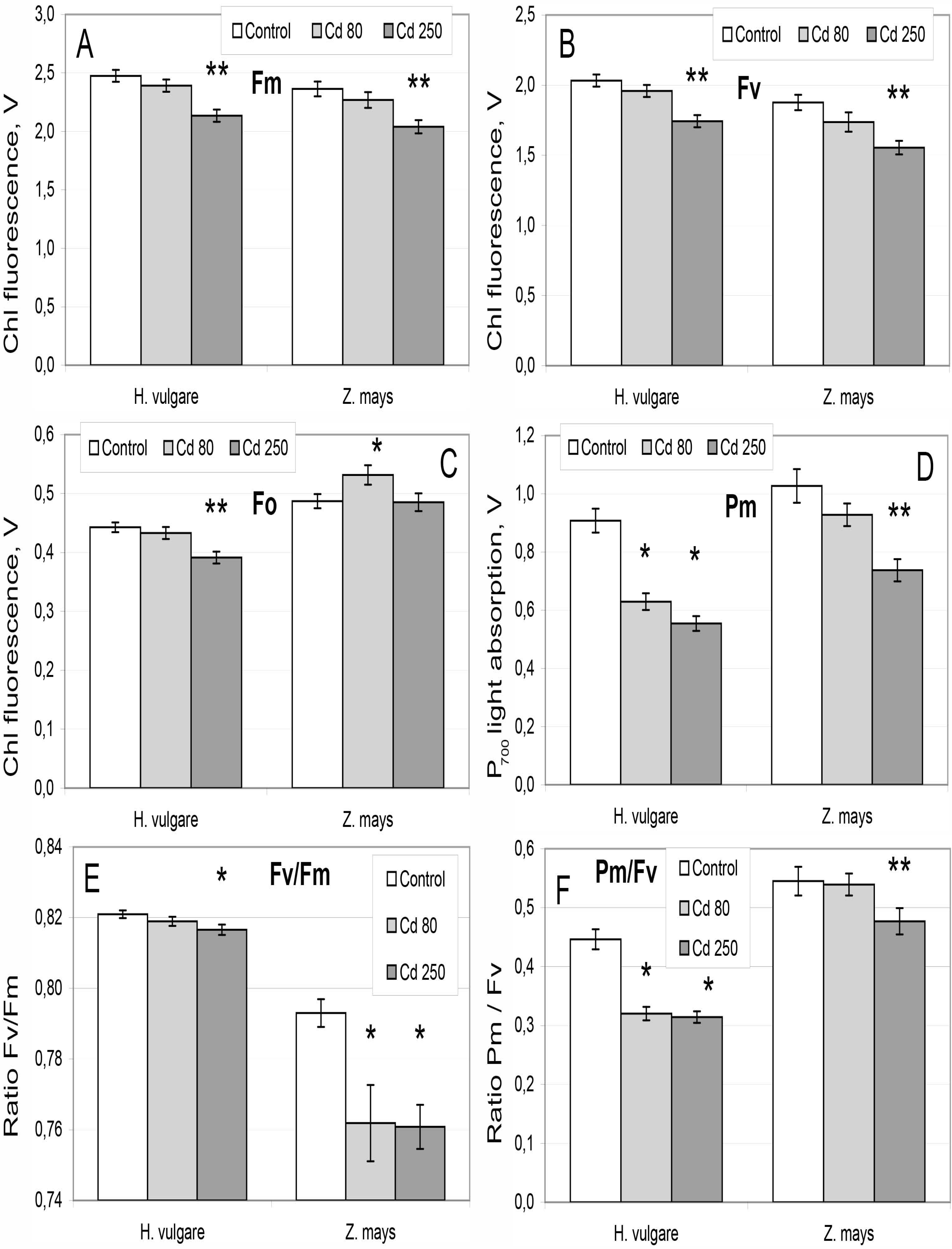
Basic photosynthetic values in the dark-adapted plants. A – Fm; B – Fv; C – Fo; D – Pm; E – Fv/Fm; F – Pm/Fv. Bars represent the variants of treatment: white – control, light grey – Cd 80 μM, dark grey – Cd 250 μM. Data are presented as means ± standard error (SE). * - the difference from control is significant at p ≤ 0.05. ** - the difference from both control and Cd 80 μM is significant at p ≤ 0.05.

Maize plants demonstrated a gradual decline of Fm, Fv, and Pm (Fig. 1); it was in accordance with both general toxicity of Cd and its accumulation in maize chloroplasts (Klaus et al., 2013; Lysenko et al., 2015). At 250 μM Cd, the ratio Pm/Fv showed larger decline of Pm comparing with Fv (Fig. 1F). The minimal Chl fluorescence demonstrated unusual changes. At 80 μM Cd, Fo level was increased; at 250 μM Cd, the level of Fo returned to the control level (Fig. 1C). The changes of Fo altered Fv/Fm dynamics. In maize, Fv/Fm dropped at 80 μM Cd with no further decrease at 250 μM Cd (Fig. 1E).

### 3.2. Chl fluorescence under light conditions

The dynamics of photochemical quenching of light energy by PSII are shown in Fig. 2. The time immediately after light induction (0.5 s) is the only point at which changes of PSII activity were proportional to Cd accumulation in the chloroplasts of both species. At this point, both 80 μM Cd and 250 μM Cd induced an equal drop of photochemical quenching in barley; the maize plants demonstrated a gradual decrease of photochemical quenching that was significant at 250 μM Cd only. The values of X(II) after 0.5 s AL are shown in Suppl. Fig. S1C with a higher resolution; the values of qP and Փ_PSII_ demonstrated the same tendency at this point.

**Fig. 2.**
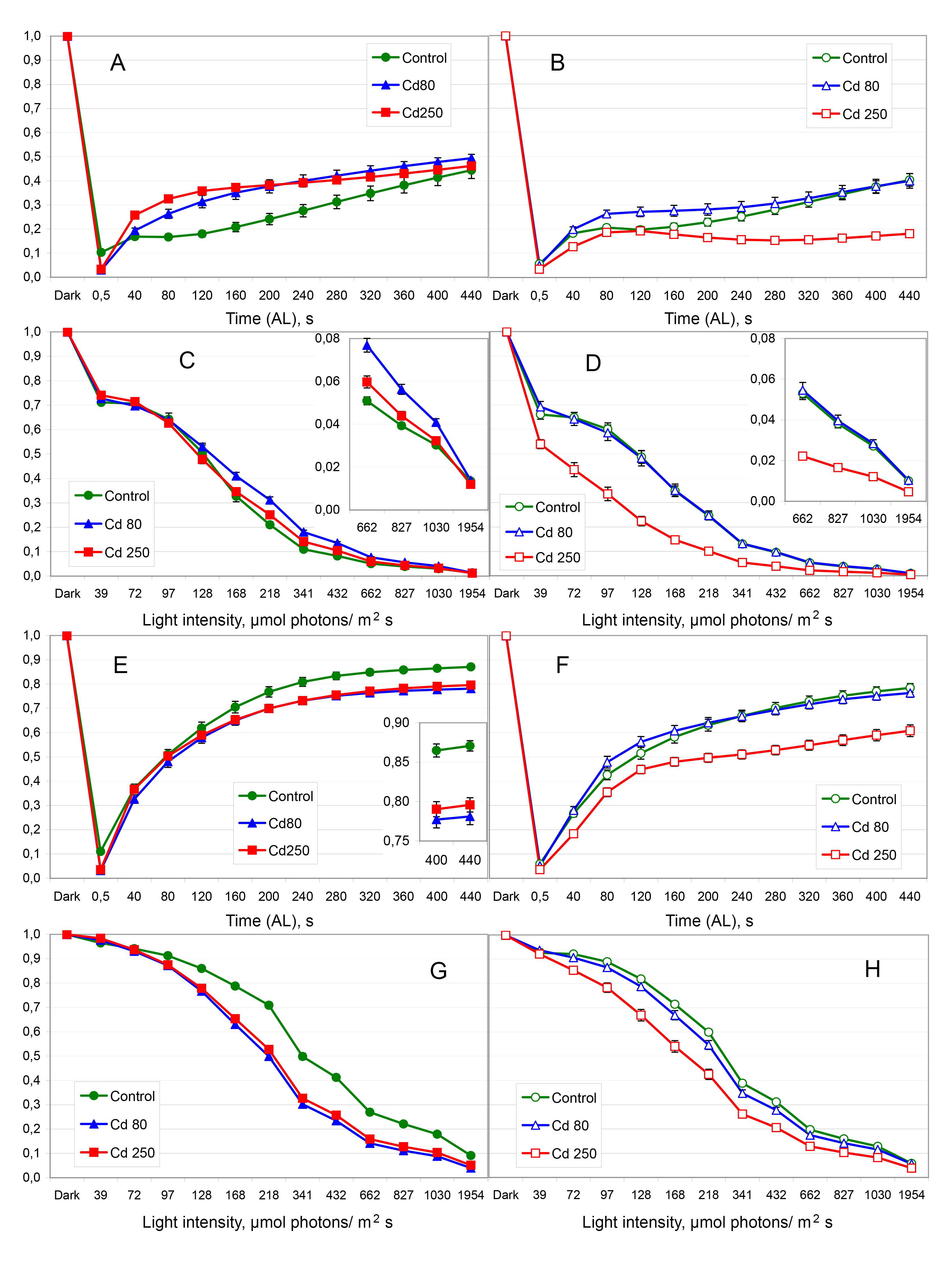
Dynamics of photochemical coefficients X(II) and qP of PSII. A-D – X(II); E-H – qP (dynamics of Փ_PSII_ are represented in Suppl. Fig. S2). A, B, E, F – IC; C, D, G, H – RLC. Lines represent the variants of treatment: green circles – control, blue triangles – Cd 80 μM, red squares – Cd 250 μM. Filled symbols represent barley; open symbols represent maize. Data are presented as means ± SE. The insets show the corresponding values with the higher resolution. “Light” denotes for AL.

The coefficient X(II) indicates the portion of photochemically active PSII in current light conditions (Fm' – Fs) comparing with the maximal level (Fv). In barley plants, Cd treatment did not reduce the portion of photochemically active PSII (Fig. 2A,C). The photochemical coefficient qP shows the balance between open (Fm' – Fs) and closed (Fs – Fo') PSII in current light conditions. The cadmium treatment significantly decreased the level of qP (Fig. 2E,G); however, it was achieved by virtue of increasing the portion of closed PSII (qC, see this section below) that was proportionally larger than the increase of open PSII. The coefficient Փ_PSII_ demonstrated an intermediate tendency (Suppl. Fig. S2A,C); the value of Փ_PSII_ is also influenced with the changes of (Fs – Fo') and Fo'.

In maize, the photochemical activity of PSII varied in accordance with Cd accumulation in the chloroplasts (Lysenko et al., 2015). At 80 μM Cd, the photochemical quenching was unchanged or slightly altered. At 250 μM Cd, the photochemical quenching decreased substantially. All the coefficients X(II) (Fig. 2B,D), qP (Fig. 2F,H), and Փ_PSII_ (Suppl. Fig. S2B,D) showed the same tendency.

Non-photochemical energy quenching is the major way reducing the photochemical quenching of PSII. In barley plants, Cd treatment reduced non-photochemical quenching (Fig 3A,C). According to IC data, both 80 μM and 250 μM Cd decreased non-photochemical quenching similarly; at the final point (440 s) solely, the effect of 80 μM Cd was significantly larger than the effect of 250 μM Cd (Fig. 3A). However, at the end of IC, all the three variants showed similar values. The application of RLC method revealed clearly that at 80 μM Cd the decrease was larger than at 250 μM Cd (Fig. 3C). Both coefficients qN (Fig. 3A,C) and NPQ (Suppl. Fig. S3A,C) demonstrated the same tendencies.

**Fig. 3.**
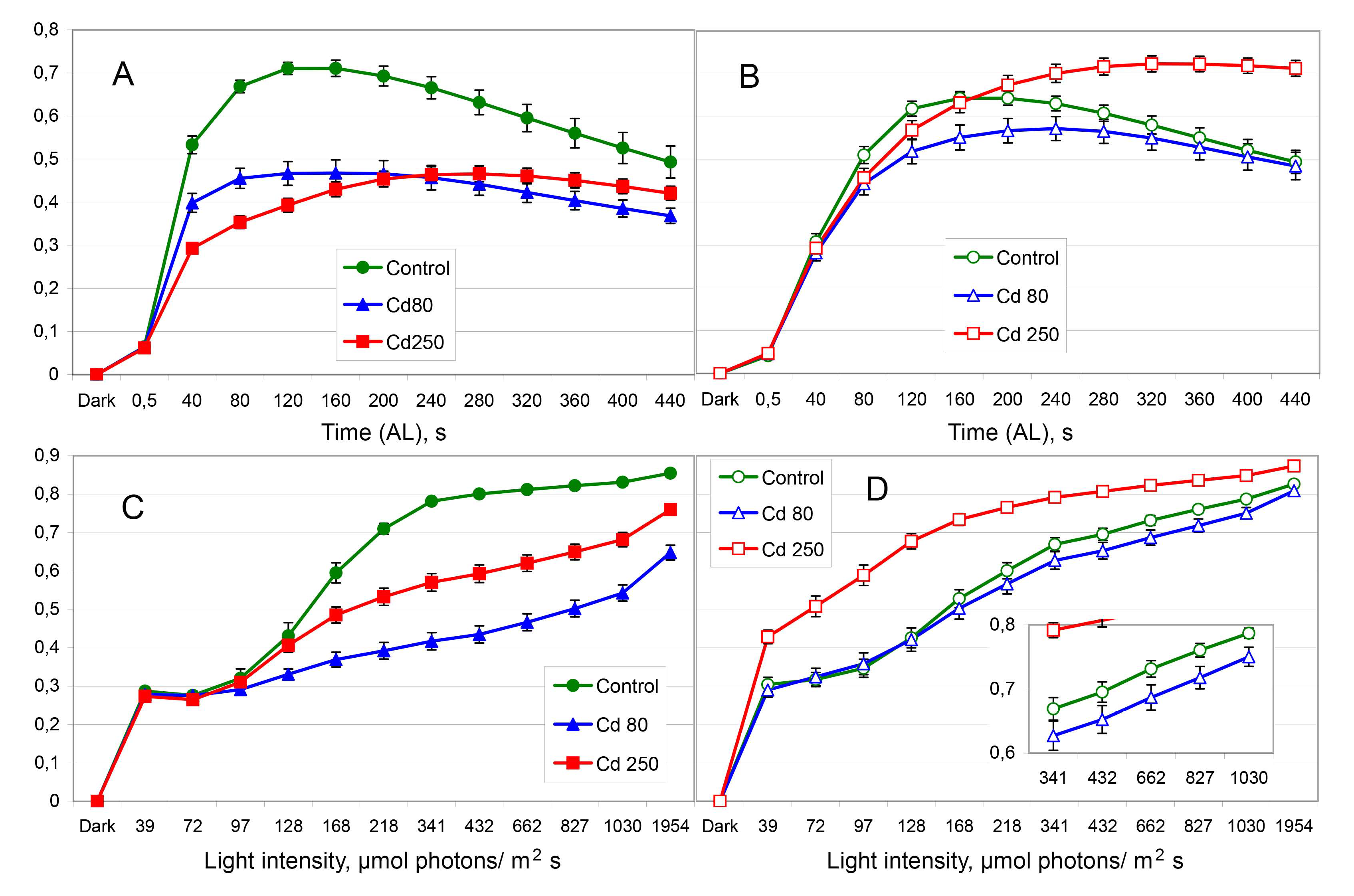
Dynamics of non-photochemical quenching (qN) of PSII. A, B – IC; C, D – RLC. All other designations are the same as in Fig. 2. Dynamics of NPQ are represented in Suppl. Fig. S3.

At 80 μM Cd, the low Cd content in maize chloroplasts (Lysenko et al., 2015) was followed by no or small decrease of non-photochemical quenching (Fig 3B,D). The small significant decreases were found in the middle of IC (80-200 s, Fig. 3B; Suppl. Fig. S3B) and at the end of RLC (AL 827-1030 μmol photons m^-2^ s^-1^, Fig. 3B; AL 662-1954 μmol photons m^-2^ s^-1^ Suppl. Fig. S3B). At 250 μM Cd, maize chloroplasts accumulated more Cd (Lysenko et al., 2015) that increased non-photochemical quenching (Fig. 3B,D; Suppl. Fig. S3B,D). Thus, Cd treatment induced the opposite changes of non-photochemical quenching in barley and maize.

A light energy that was neither photochemically nor non-photochemically quenched, is retained in so called closed PSII complexes. The portion of closed PSII, calculated using the coefficient qC, is represented in Fig. 4. In barley plants, Cd treatment increased the portion of closed PSII. The method IC demonstrated relatively small and equal effect of both 80 μM and 250 μM Cd (Fig. 4A). Applying RLC method, we revealed a high level of the closed PSII accumulation at 80 μM Cd; at 250 μM Cd, the effect was smaller, while yet rather large (Fig. 4C).

**Fig. 4.**
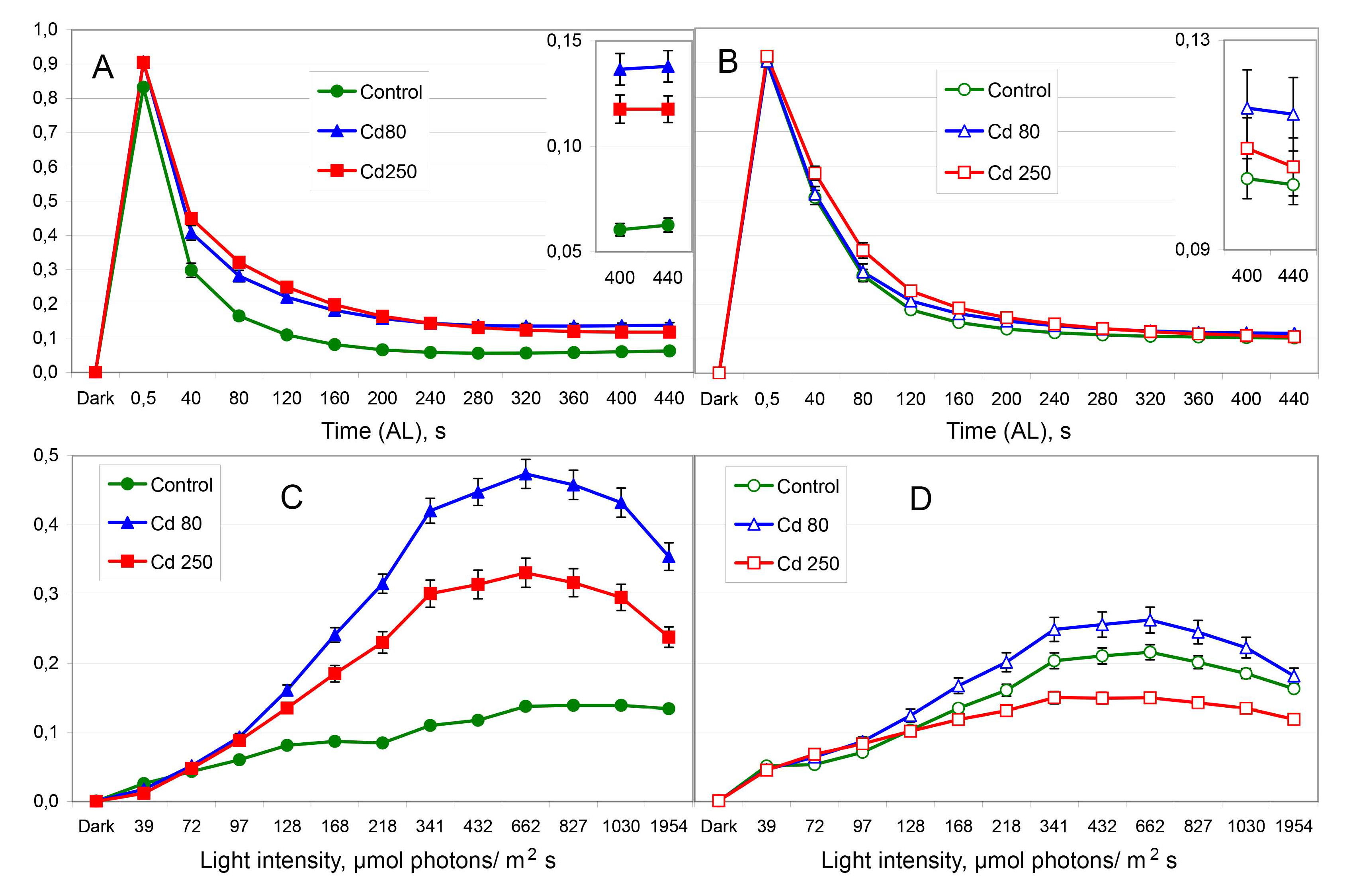
Dynamics of closed PSII (qC). A, B – IC; C, D – RLC. All other designations are the same as in Fig. 2.

In maize plants, Cd imposed little effect on qC dynamics. The method IC demonstrated no influence of Cd treatment (Fig. 4B). The method RLC revealed small bidirectional changes of qC (Fig. 4D). At 80 μM Cd, the portion of closed PSII was above the control level; at 250 μM Cd, the portion of closed PSII was below the control level. Thus, RLC dynamics revealed the similarity between barley and maize. In both species, elevating the concentration from 80 μM Cd to 250 μM Cd stimulated the increase of qN (Fig. 3B,D) and decrease of qC (Fig. 4B,D).

### 3.3. P_700_ light absorption under light conditions

In the dark, the level of Y(I) in the control plants was substantial. In maize, it was two times larger than in barley (Fig. 5A,B); it is shown with a higher resolution in Suppl. Fig. S1A. This assumes the higher level of cyclic electron transport in maize. In maize plants, Cd treatment gradually decreased Y(I) in darkness; in barley plants, Cd treatment increased Y(I) in darkness (Suppl. Fig. S1A).

**Fig. 5.**
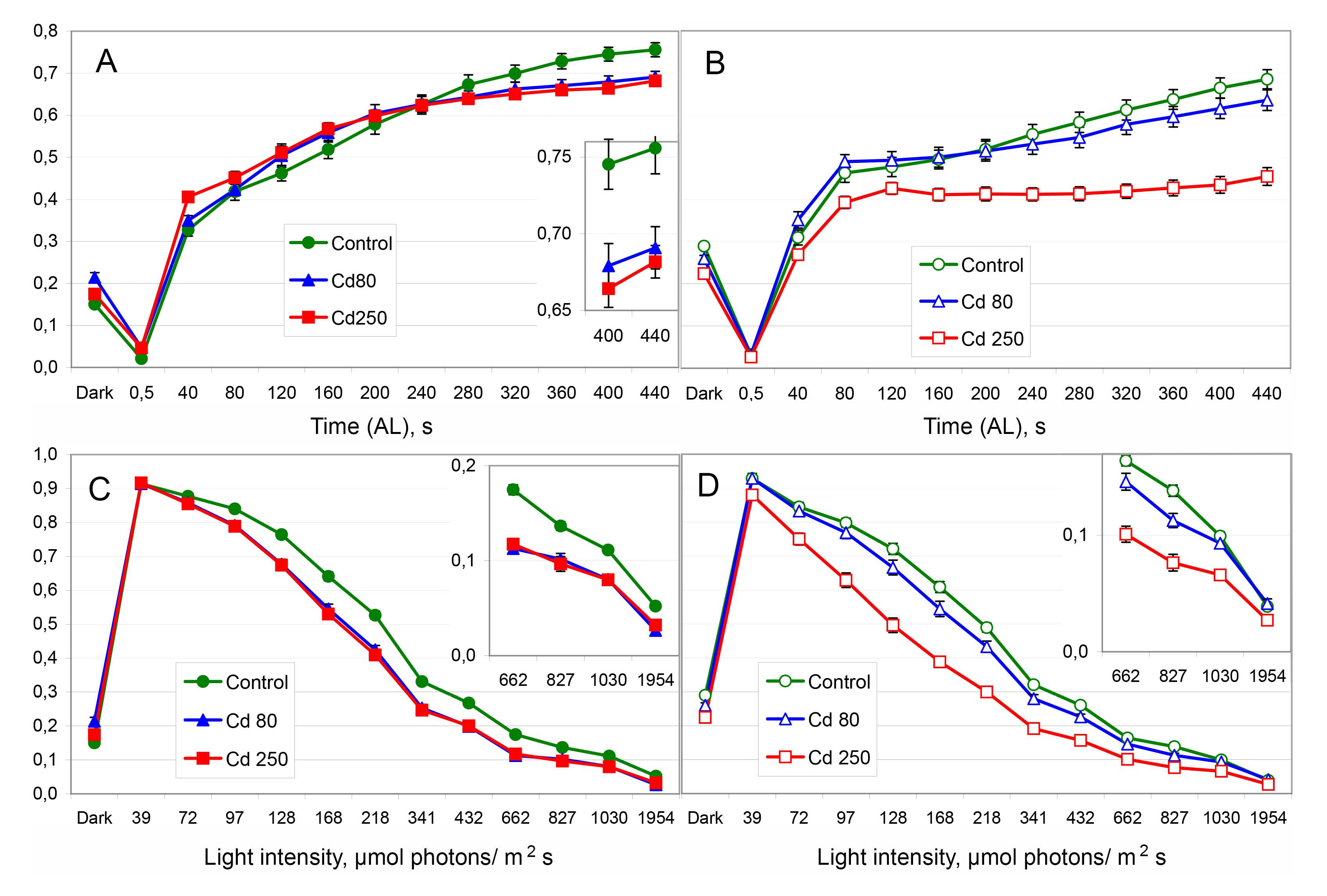
Dynamics of the quantum yield Y(I) of PSI. A, B – IC; C, D – RLC. All other designations are the same as in Fig. 2.

Under illumination, the photochemical activity of PSI showed a reasonable correlation with Cd accumulation in the chloroplasts. In barley plants, 80 μM Cd and 250 μM Cd induced the same decrease of Y(I) (Fig. 5A,C); the differences from the control were significant in IC from 360/320 s and in RLC from AL 97/72 μmol photons m^-2^ s^-1^ (for the concentrations 80/250 μM Cd respectively). In maize plants, 80 μM Cd imposed no (IC) or very small (RLC) decrease of Y(I) (significant at AL 168-827 μmol photons m^-2^ s^-1^; p=0.0504 at AL 128 μmol photons m^-2^ s^-1^). At 250 μM Cd, we revealed a large reduction of Y(I) in maize (Fig. 5B,D).

The photochemical activity of PSI was limited mostly at the donor side (Fig. 6). Donor-side limitation Y(ND) reached its maximal level in 1-2 min after the light induction and then declined slowly in the most cases (Fig. 6A,B). In barley plants, Cd treatment accelerated this process; the same maximal level of Y(ND) was reached and began to decline 40 s earlier than in the control (Fig. 6A). However, Y(ND) relaxed slower in Cd treated plants than in the control; at the end of IC, the near stationary level of Y(ND) was increased in Cd treated plants (significant from 360 or 400 s). In RLC, Y(ND) was increased in all the range of AL intensities; the increase was significant in all but one point (AL 1954 μmol photons m^-2^ s^-1^ at 80 μM Cd) (Fig. 6C). The effect of 80 μM and 250 μM Cd was equal (Fig. 6A,C). In maize plants, 80 μM Cd induced a minor significant increase of Y(ND) while 250 μM Cd induced a significantly larger increase of Y(ND); the differences were significant at all points of both IC and RLC (Fig. 6B,D).

**Fig. 6.**
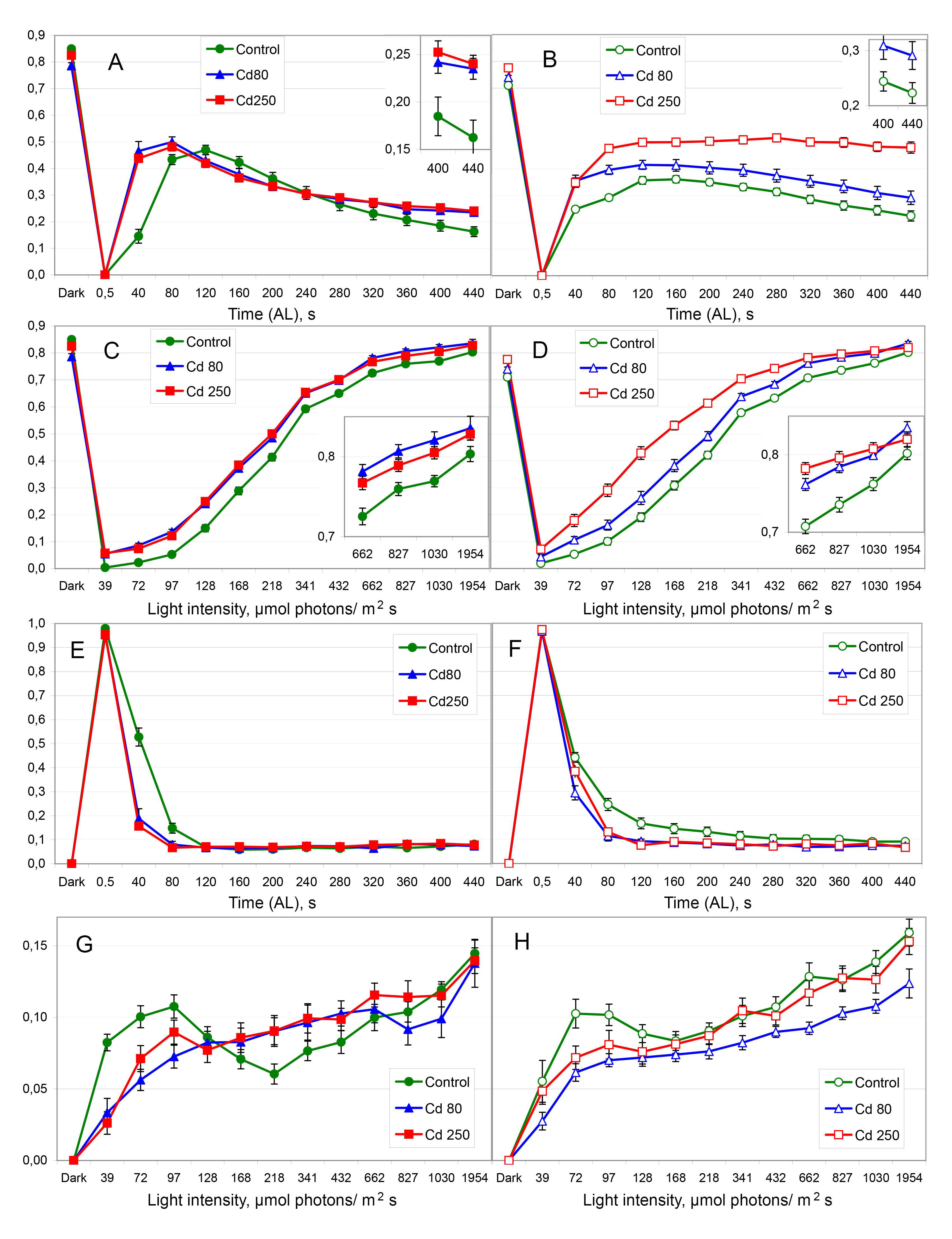
Dynamics of the limitations of PSI. A-D – limitation at the donor side Y(ND); E-H – limitation at the acceptor side Y(NA). A, B, E, F – IC; C, D, G, H – RLC. All other designations are the same as in Fig. 2.

The impact of acceptor-side limitation Y(NA) was small (Fig. 6E-H), while it was increased slowly along with the increase of AL intensity (Fig. 6G,H). The sole exception was the immediate reaction to the light induction; at 0.5 s after the light induction, the photochemical activity of PSI was restricted mostly at the acceptor side (Fig. 6E,F). At this time, the photochemical activity of PSI Y(I) demonstrated no correlation with Cd accumulation in the chloroplasts (Fig. 5A,B); this time slot is shown with a higher resolution in Suppl. Fig. S1B.

### 3.4. Ratios of PSII and PSI activities

The photochemical activities of PSI and PSII can be compared with the use of ratio Y(I)/X(II) (Lysenko et al., 2020). In the control, both plant species demonstrated similar dynamics of Y(I)/X(II) in IC (Fig. 7A,B) and RLC (Fig.7C,D; Suppl. Fig.S5). In barley, Cd treatment decreased the ratio Y(I)/X(II); this means that the balance of photosystems was shifted in favor of PSII. The effect of 250 μM Cd was smaller than the effect of 80 μM Cd (Fig. 7A,C). The effect of Cd was dissimilar to the effects of other heavy metals, namely Cu and Fe (Fig. 7A). In maize, the treatment with 80 μM Cd imposed no or small decrease of Y(I)/X(II) (80-200 s in IC, AL 662-827 μmol photons m^-2^ s^-1^ in RLC). On the contrary, the treatment with 250 μM Cd increased the ratio Y(I)/X(II) (Fig. 7B,D); this increase will be discussed in section 4.4.

**Fig. 7.**
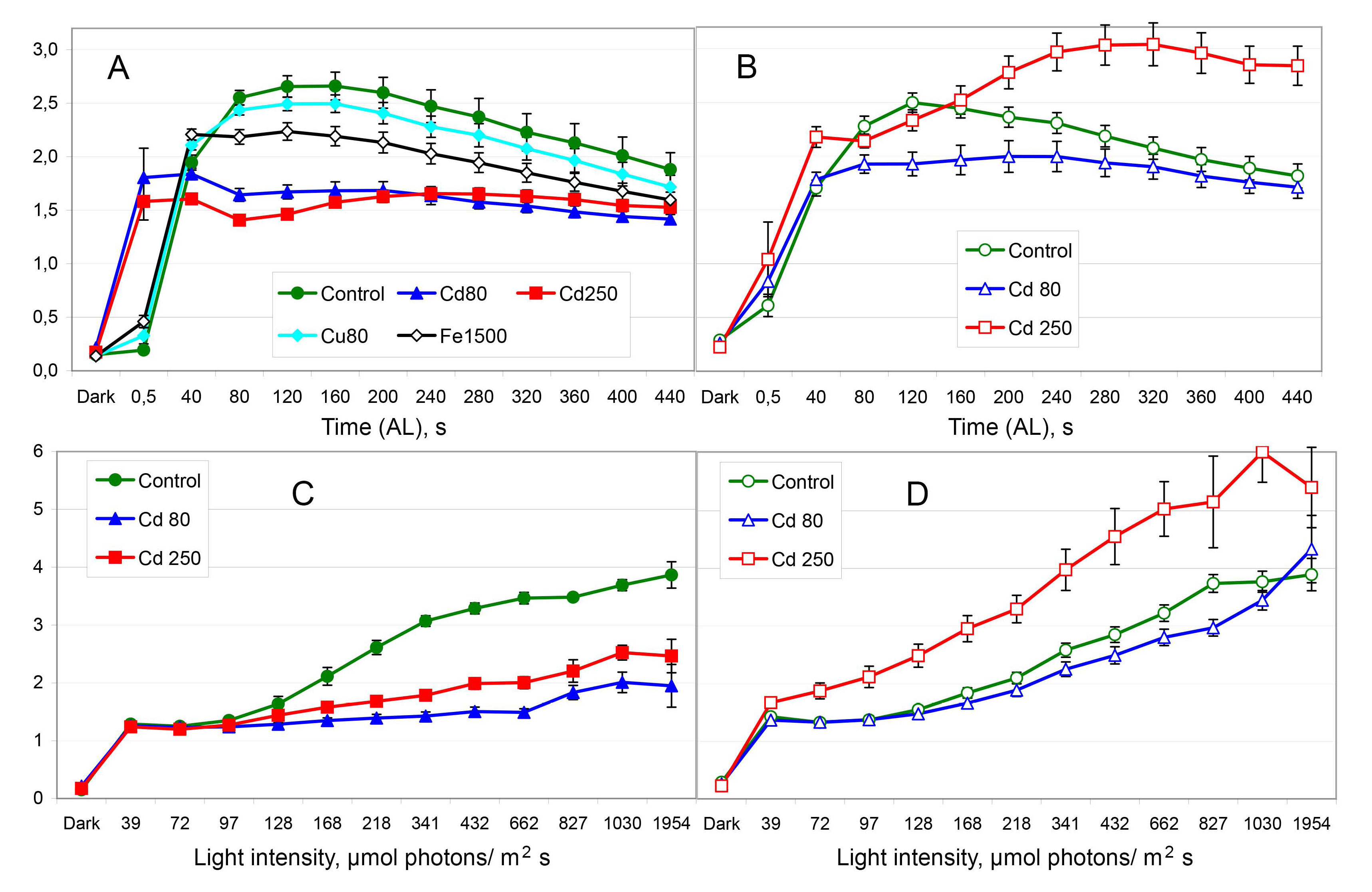
The ratio Y(I)/X(II) showing balance between the photochemical activities of PSI and PSII. A, B – IC; C, D – RLC. Lines with diamond symbols (A) represent additional variants of treatment: turquoise filled – Cu 80 μM, black open – Fe 1500 μM. All other designations are the same as in Fig. 2.

The coefficients qC, Y(ND), and Y(NA) are calculated similarly; each of them shows which part of the maximal ability to photochemical quenching (Fv or Pm) was lost due to particular limitation. The ratio qC/Y(NA) enables comparing limitations at the acceptor-sides of PSII and PSI correspondingly. Actually, the acceptor side of PSI is limited mostly by the activity of Calvin-Benson cycle; the acceptor side of PSII is limited by the most part of the electron-transfer chain. So, the ratio qC/Y(NA) enables comparing limitations of the electron-transfer chain and Calvin-Benson cycle. Some data points were excluded from the analysis. In darkness, Y(NA) is zero and qC is close to zero that makes the analysis impossible and meaningless. In RLC at the first AL intensity (39 μmol photons m^-2^ s^-1^), many Y(NA) data were small negative, zero, or very small positive; therefore, too large portion of data points should be discarded (see section 2.3) that makes correctness of the results questionable.

Immediately (0.5 s) after the light induction, both acceptor-side limitations were high and their ratio was close to 1 (Fig. 8A,B). In barley, the acceptor-side limitation of PSI exceeded that of PSII; it was observed in the control plants (qC/Y(NA) = 0.85) and also in Cu-and Fe-treated plants (qC/Y(NA) = 0.87-0.88). Cadmium treatment equalized both limitations (qC/Y(NA) = 0.95). In maize, all the variants were equal (qC/Y(NA) = 0.93-0.95) (Fig. 8A,B insets). The ratio reached the peak and then declined. In Cd-treated plants the peak was reached in 80 s; in maize at 250 μM Cd, the peak was reached in 80 or 120 s. In control barley plants, the peak was reached later (120 s); control maize plants demonstrated no clear peak (Fig. 8A,B). Under illumination 128 μmol photons m^-2^ s^-1^, control plants reached the steady state level of about 1 that means an equality of both limitations. Cadmium treatment increased qC/Y(NA) values; the steady state level was close to 2 (Fig. 8A,B). The latter means that qC was two times larger than Y(NA). In RLC at the diverse AL intensities, qC/Y(NA) was >1 usually; however, control plants demonstrated low values qC/Y(NA) at low AL intensities (Fig. 8C,D). Generally, qC/Y(NA) dynamics mirrored the dynamics of qC (Fig. 4); probably, the only exception was the time slot 0.5 s since the light induction.

**Fig. 8.**
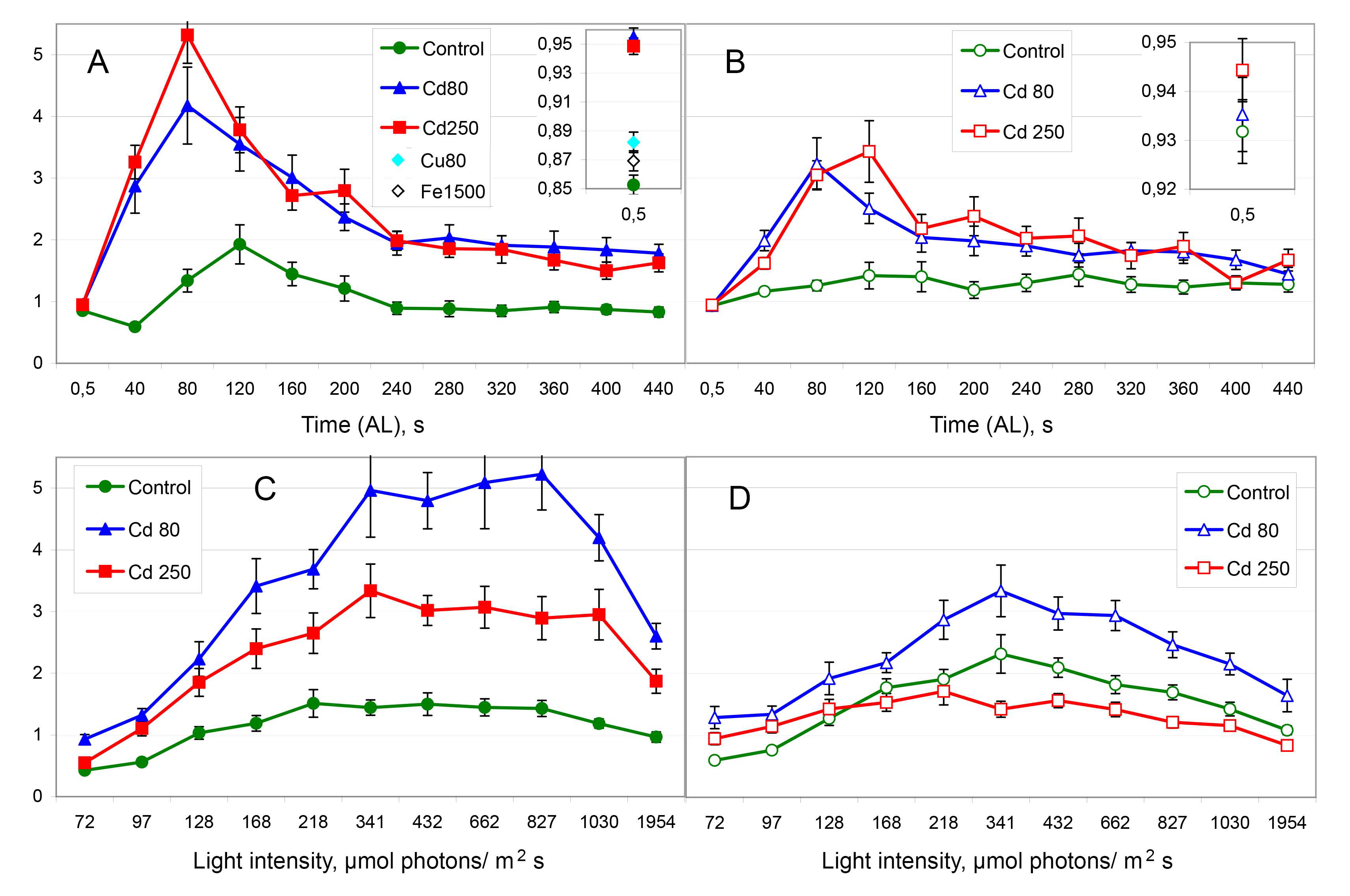
The ratio qC/Y(NA) comparing acceptor-side limitations of PSII (qC) and PSI (Y(NA)). A, B – IC; C, D – RLC. All other designations are the same as in Fig. 2 and Fig. 7.

The ratio qC/Y(ND) enables comparing limitations in the section of electron-transport chain between the acceptor side of PSII and donor side of PSI. Some data points were excluded from the analysis. In darkness, qC is nearly zero that makes the analysis senseless. Immediately (0.5 s) after the light induction, Y(ND) is zero that makes the analysis impossible and senseless. In the beginning of RLC, control plants and also Cu-and Fe-treated plants (not shown) demonstrated too large (at 39 μmol photons m^-2^ s^-1^) or large (at 72 μmol photons m^-2^ s^-1^) portion of small negative, zero, or very small positive Y(NA) data that should be discarded (see section 2.3); it makes correctness of the results questionable. We hesitated, however, excluded the point AL 39 μmol photons m^-2^ s^-1^ from the analysis and included the point AL 72 μmol photons m^-2^ s^-1^ to the analysis.

In the beginning of IC, the ratio qC/Y(ND) was able to exceed 1; control barley plants also demonstrated qC/Y(ND) > 1 in the beginning of RLC. In these cases, the limitation at the acceptor side of PSII (qC) prevailed. The most parts of the dynamics showed the ratio qC/Y(ND) < 1; this means that limitation at the acceptor side of PSII (qC) was mostly smaller than limitation at the donor side of PSI (Y(ND)). These data are shown in Suppl. Fig. S6. In barley, Cd treatment enlarged the ratios qC/Y(ND) (Suppl. Fig. S6A,C). In maize, 80 μM Cd had a limited decreasing effect on the ratio; 250 μM Cd decreased qC/Y(ND) obviously (Suppl. Fig. S6B,D).

The data set shown in Suppl. Fig. S6 was revisualized and presented in Fig. 9. Both IC and RLC of barley and maize were gathered to show all the dynamics in the same space. For better visibility, the data were restricted with qC/Y(ND) values from 0 to 1 and divided into three panels. Fig. 9 shows that all the dynamics decreased to a quasi-stationary level that was kept for 1-2 min. In a variant (species/Cd concentration), this level was the same in both IC and RLC; value of the level varied. Control barley plants and maize plants treated with 250 μM Cd demonstrated the level 0.2 meaning that qC was five times lower than Y(ND) (Fig. 9A). Maize plants in control and under 80 μM Cd showed the level 0.4 meaning that qC was 2.5 times smaller than Y(ND) (Fig. 9B). Cadmium treated barley plants were stabilized shortly at the level 0.5 meaning that qC was two times lower than Y(ND) (Fig. 9C). The only exception was RLC of barley treated with 80 μM Cd keeping the level 0.65 meaning that qC was reduced by one third comparing with Y(ND; the corresponding IC was stabilized at the level 0.5 (Fig. 9C).

**Fig. 9.**
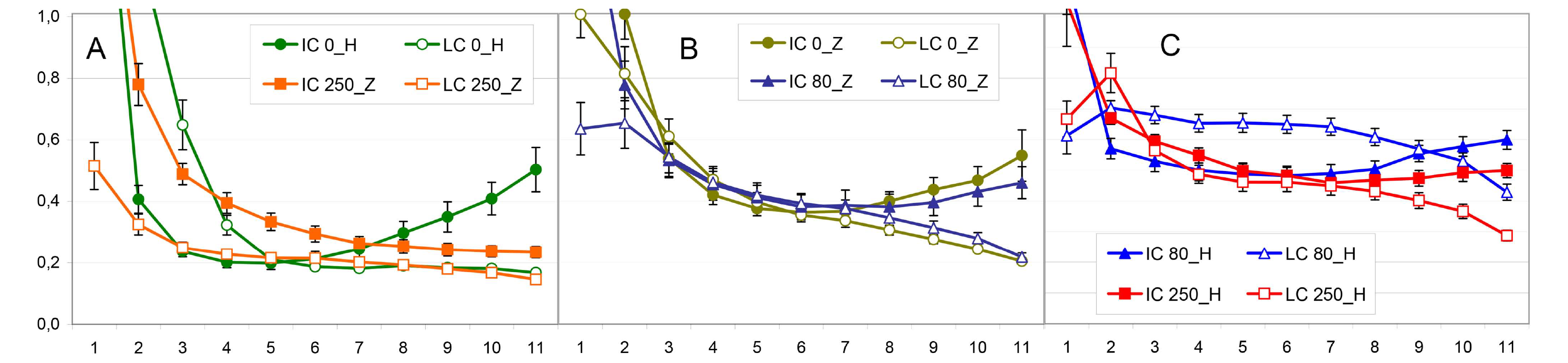
The ratio qC/Y(ND) showing balance of limitations between the acceptor side of PSII (qC) and the donor side of PSI (Y(ND)). The figure revisualizes the complete data set presented in Suppl. Fig. S6. For the better visibility, the values >1 are omitted, and the curves are separated into panels A-C according to sizes of the quasi-stationary levels (see Sections 3.4 and 4.5). The symbols represent: filled – IC and open – (R)LC; circles – control, triangles – Cd 80 μM, squares – Cd 250 μM. The shades of green mark the control: bright – barley and brownish – maize. The shades of blue denote Cd 80 μM treatment: bright – barley and dark – maize. The shades of red denote Cd 250 μM treatment: red – barley and orange – maize. H – *H. vulgare*; Z – *Z. mays*; 0, 80, and 250 - μM Cd added. The abscissa numbering indicates both time and AL intensities. For IC, 1-11 correspond to the range 40 - 440 s; for RLC, 1-11 correspond to the range 72 – 1954 μmol photons m^-2^ s^-1^. Data are presented as means ± SE.

In IC, a short period of stabilization was followed by an increase in qC/Y(ND) ratio. The increase was larger in barley than in maize; the increase became less pronounced with the rise of Cd concentration; at 250 μM Cd, maize demonstrated no such increase (Fig. 9). In RLC, the intensity of AL was magnified stepwise; therefore, a stabilization was followed by a decrease of qC/Y(ND) ratio. The higher was the quasi-stationary level, the larger was the subsequent decrease (Fig. 9). In the control, the short-time stabilization level in barley (Fig. 9A) was lower than that in maize (Fig. 9B). In barley, Cd treatment increased the stabilization level (Fig. 9A,C).

In maize, Cd treatment decreased the stabilization level (Fig. 9A,B). These data suggest an ability of the photosynthetic apparatus keeping a connection between changes at the acceptor side of PSII and donor side of PSI.

## 4. Discussion

We used the model of two Poaceae species with different pattern of Cd accumulation in the chloroplasts (Lysenko et al., 2015). We applied this model to distinguish between direct and indirect effects of Cd on the processes in the electron-transport chain in chloroplasts. An indirect effect should reflect the general toxicity and be increased along with the increase of Cd concentration in hydroponics from 80 μM to 250 μM; this increase should be similar in both species (Klaus et al., 2013; Lysenko et al., 2015). A direct effect should reflect Cd content in the chloroplasts (Lysenko et al., 2015). At 80 μM Cd, a direct effect should be small in maize and large(r) in barley. At the higher concentration 250 μM Cd, a direct effect should be increased in maize and showed no increase in barley.

### 4.1. Maximal parameters in darkness

The adaptation to darkness is applied to reduce any limitation of photochemical energy quenching by PSII and PSI; the parameters measured in the dark-adapted state reflect quantities of functionally active reaction centers of PSII (Chl fluorescence) and PSI (P_700_ light absorption). The level of Pm depends on the quantity of reaction centers of PSI and light scattering (Dual-PAM_1e, 2009).

The parameters of Chl fluorescence (Fm, Fv, Fo, Fv/Fm) demonstrated a gradual decline along with the growth of Cd content in mineral media (Fig. 1). In maize, Fo changed nonlinearly (Fig. 1C), which correspondingly influenced Fv/Fm alteration (Fig. 1E).

The maximal change of P_7 00_ (Pm) altered in agreement with Cd content in the chloroplasts (Fig. 1D). Probably, the gradual slope of Fm and Fv reflected just the reduction of leaf size and/or chloroplast density in these leaves. So, normalized Pm (Pm/Fv) showed better a correlation with Cd content in the chloroplasts. In barley, Pm/Fv significantly reduced at 80 μM Cd with no more reduction at 250 μM Cd; in maize, Pm/Fv remained unchanged at 80 μM Cd and decreased at 250 μM Cd (Fig. 1F).

The larger decrease in Pm than in Fv was rather specific to Cd stress. Cu treatment decreased Pm/Fv to a lesser extent, Fe treatment unchanged it (Lysenko et al., 2020), and heat stress increased Pm/Fv in barley and maize plants of the same age (Lysenko et al., 2023).

These data suggest that the quantity of functionally active PSII was diminished with an indirect action of Cd while the reduction of functionally active PSI can be explained by a direct Cd action in the chloroplasts. The changes of Fo and Fv/Fm in maize can be explained neither by an indirect nor by a direct Cd action; they probably resulted from a simultaneous action of direct and indirect mechanisms that can not be unraveled within the framework of the current experimental model.

### 4.2. PSI activity under illumination

The dynamics of P_700_ parameters changed in agreement with Cd accumulation in the chloroplasts (Lysenko et al., 2015). In barley, the quantum yield of PSI Y(I) decreased moderately at 80 μM with no further reduction at 250 μM (Fig. 5A,C). In maize, Y(I) decreased slightly at 80 μM Cd and showed a large decrease at 250 μM Cd (Fig. 5B,D).

The decrease of PSI photochemical quenching was caused by an increasing limitation at the donor side (Y(ND)). In barley, the increase of Y(ND) was equal at both 80 μM and 250 μM Cd; in Cd-treated barley plants, Y(ND) remained higher at the end of IC and all over RLC (Fig. 6A,C). In maize, 80 μM Cd stimulated a small increase of Y(ND) while 250 μM Cd induced a large increase of Y(ND) (Fig. 6B,D); the control, 80 μM Cd, and 250 μM Cd differed significantly from each other throughout both IC and RLC. The effect of Y(ND) increasing was rather specific to Cd. The excess of Cu also increased Y(ND) while the excess of Fe did not increase Y(ND) (Lysenko et al., 2020). The heat stress decreased Y(ND) in nine-day-old barley and maize plants (Lysenko, unpublished data).

Limitation of PSI at the acceptor side Y(NA) was small and showed no correlation with Cd content in the chloroplasts (Fig. 6E-H). Each Cd treatment decreased Y(NA) rapidly at the beginning of IC (Fig. 6E,F). In RLC, control plants showed sigmoid-like dynamics; at the extremum points, they differed significantly from more linear dynamics of Cd-treated plants. The only case of the overall Y(NA) decrease was observed in RLC of maize plants treated with 80 μM Cd (Fig. 6F); in this case, Cd content in the chloroplasts was minimal (Lysenko et al., 2015).

Thus, we observed the relationship between Cd content in chloroplasts and Cd effect on PSI. At 80 μM Cd, maize chloroplasts accumulated a small amount of Cd (49 ± 5 ng/mg Chl); in this condition, PSI activity was not influenced at all (Pm/Fv) or significantly (Pm, Y(I)) whereas its limitation Y(ND) was slightly increased. At 250 μM Cd, maize chloroplasts accumulated more Cd (92 ± 20 ng/mg Chl) (Lysenko et al., 2015); this magnification was significant at p = 0.0754 only. However, it was followed by a rather large significant inhibition of PSI (Pm, Pm/Fv, Y(I)) and an increasing acceptor-side limitation (Y(ND)) of PSI. More probably, Cd content was increased in the maize chloroplasts at 250 μM Cd comparing with 80 μM Cd; yet, more than four repeats of the experiment were required to fit into p < 0.05. We observed higher accumulation of Cd in the first and second leaves of maize at 250 μM Cd than at 80 μM Cd; however, each time the difference was insignificant. It was observed in two independent studies with the same experimental design (Klaus et al., 2013; Lysenko et al., 2015) and with a prolonged treatment (Klaus et al., 2013); in the latter case, the difference was significant for the third leaves solely (leaves of upper storeys accumulated Cd equally). Under 250 μM Cd, maize plants were severely damaged and 42% of them died in a one-month treatment. This increased their biological variability; in some cases, we had to repeat simple experiments with leaves 10-12 times (Klaus et al., 2013).

At 80 μM Cd, barley chloroplasts accumulated a large amount of Cd (171 ± 26 ng/mg Chl). At 250 μM Cd, barley chloroplasts accumulated less Cd (126 ± 7 ng/mg Chl); the decrease was insignificant (Lysenko et al., 2015). At both 80 μM and 250 μM Cd, the activity of PSI decreased equally (Pm/Fv, Y(I)) or the difference was insignificant (Pm); donor-side limitation Y(ND) was increased equally as well. Up to this, we considered that both Cd content in the chloroplasts and Cd effect on PSI was not increased at 250 μM comparing with 80 μM Cd; it is true. However, whether the equal effect on PSI was induced by equal or even smaller content of Cd at 250 μM comparing with 80 μM Cd?

There are several possible explanations. First, we can assume both contents equal using insignificance of the difference as the key reason. But it does not seem reliable. In the leaves of barley plants, Cd content decreased at 250 μM comparing with 80 μM Cd; the decrease was significant and substantial (Lysenko et al., 2015). The decrease of Cd content in the leaves practically unchanged the proportion of Cd introducing into the chloroplasts. The current experimental design with 80 μM Cd and barley is the advantageous model; we used it in a series of studies. At 80 μM Cd, barley chloroplasts accumulated 237 ± 30 ng Cd/mg Chl (Lysenko et al., 2019) and 214 ± 22 ng Cd/mg Chl (Lysenko et al., 2020); both values were significantly higher than 126 ± 7 ng Cd /mg Chl accumulated at 250 μM Cd (Lysenko et al., 2015). One more treatment was one day shorter; eight-day-old barley plants accumulated in the chloroplasts 152 ± 6 ng Cd /mg Chl (N=10) (Lysenko et al., unpublished data) that was also significantly higher than 126 ± 7 ng Cd /mg Chl. In one more study, barley chloroplasts accumulated the same amount of Cd (125 ± 6 ng/mg Chl, N=8) (Lysenko et al., unpublished data); however, one parameter of experimental design was changed in this experiment that can influence Cd accumulation. More probably, barley plants accumulated less Cd in both leaves and chloroplasts at 250 μM Cd than at 80 μM Cd.

Second, we can admit that though Cd contents in the chloroplasts were different, but the difference was too smal and/or exceeded a threshold level; therefore, the same direct effect was imposed both at 80 μM and at 250 μM Cd. Third, we can suppose that Cd direct effect was also smaller at 250 μM Cd; however, it was supplemented by an indirect Cd effect; therefore, the entire effects were equal at both 80 μM and 250 μM Cd. We regard the latter explanation as more probable. Both barley and maize plants were reduced much more at 250 μM comparing with 80 μM Cd; the higher general toxicity should increase numerous indirect effects in both species. In maize, a direct Cd effect on PSI could also be supplemented by an indirect effect; however, we cannot distinguish it within the framework of this experimental model. The interaction of a direct and an indirect effects is discussed in section 4.3.3 considering the changes of closed PSII reaction centers.

The analyses of PSI activity under Cd stress are scarce. *In vitro*, Cd treatment had no effect on PSI activity (Bazzaz and Govindjee, 1974). *In vivo*, Cd treatment inhibited PSI activity at the acceptor side in 21-day-old maize plants (Siedlecka and Baszynski, 1993). In barley chloroplasts, Cd was concentrated in thylakoids; the quantity of Cd was sufficient to replace Cu in plastocyanin (Lysenko et al., 2019). Cd-substituted plastocyanin is routinely used in analytical studies as a variant incapable to perform electron transfer (e.g., Díaz-Moreno et al., 2005). The hypothesis of Cd/Cu substitution in plastocyanin explains the mechanism of direct inhibition of PSI at the donor side (Fig. 1,5,6) and its correspondence with the changes of Cd content in the chloroplasts (Lysenko et al., 2015).

Two more features should be mentioned. In darkness, Y(I) was higher in maize than in barley that should be attributed to the higher impact of cyclic electron transport around PSI in C_4_-plants (Suppl. Fig. S1A). In barley, Cd treatment increased Y(I) in the dark. In maize, Cd treatment gradually decreased Y(I) in the dark; though, it remained higher than in barley. Recently, we have revealed that Y(I) in the dark was increased with a heat stress in both barley and maize (Lysenko et al., 2023). Probably, these species are able increasing an electron donation from stromal reductants to PSI (Havaux, 1996; Yamane et al., 2000; Rumeau et al., 2007); however, this ability was inhibited by Cd treatment in maize.

In barley plants, Cd stress accelerated PSI readaptation to the light conditions. Cd treated plants reached maximum (peak) and started a relaxation of Y(ND) faster than control plants (Fig. 6A). In the beginning of IC, Cd treatment decreased Y(NA) and it reached the steady state level earlier than in the control (Fig. 6E). Under tolerated heat stress (37-42℃), the quantum yield Y(I) reached the steady state level earlier than in the control (Lysenko et al., 2023). In maize plants, such acceleration was absent (Fig. 6B), smaller (Fig. 6F), or not so obvious (Lysenko et al., 2023).

### 4.3. Dynamics of Chl fluorescence under illumination

The dynamics of Chl fluorescence represented the complex interplay between the photochemical quenching (Fig. 2), non-photochemical quenching (Fig. 3), and portion of closed PSII complexes (Fig. 4) that was influenced with species specific features. The corresponding coefficients - X(II), qN, and qC – are comparable. They have the common denominator (Fv) and X(II) + qN + qC = 1, where 1 means Fv (Lysenko et al., 2020). If one of them increases, then the sum of two others will decrease and vice versa. It is advantageous beginning the analysis of PSII activity from non-photochemical quenching.

#### 4.3.1 Non-photochemical quenching

In the control, both species showed similar dynamics of non-photochemical quenching; the dynamics were very similar in IC and rather similar in RLC (Fig. 3, Suppl. Fig. S4). Under Cd stress, barley plants demonstrated the tendency decreasing non-photochemical quenching and a broader range of changes; maize plants demonstrated the tendency increasing non-photochemical quenching and a narrower range of variation (Fig. 3). The lowest qN values were found in barley; the highest qN values were observed in maize (Suppl. Fig. S4). The changes observed can be clearly referred to neither a direct nor an indirect mechanism of Cd action; a more complicated explanation is required.

At 80 μM Cd, qN was decreased (Fig. 3). The decrease was large in barley and very small in maize that corresponded to the large and small Cd contents in their chloroplasts (Lysenko et al., 2015). This decrease can be attributed to a direct Cd action. At 250 μM Cd, the effect was reversed. The level of non-photochemical quenching was increased at 250 μM Cd comparing with 80 μM Cd; it was obvious in three cases (Fig. 3B-D). At 250 μM Cd, barley plants slowed qN dynamic since the light induction; the “peak” with further decrease was reached later (Fig. 3A). Therefore, qN was significantly higher under 250 μM comparing with 80 μM Cd only at the last point of measurement (440 s). This reverse of effect at 250 μM Cd cannot be ascribed to the changes of Cd content in the chloroplasts and a direct Cd action. The reversed tendency was dissimilar in barley and maize; hence, we have no reason to suggest an indirect effect due to increasing of general toxicity. However, the changes of non-photochemical quenching (Fig. 3) were mirrored with the levels of closed PSII (Fig. 4); the latter can be explained by an indirect action of Cd (see section 4.3.3). Perhaps, a direct decreasing effect at 80 μM Cd was overcome by an indirect increasing Cd action on non-photochemical quenching under 250 μM Cd.

Cd treatment can change non-photochemical quenching both up and down. It was observed in different studies (Burzynski and Zurek, 2007; He et al., 2008; Lysenko et al., 2015; 2020) and even in a single experiment. The opposite Cd effect was obtained at different concentrations: qN was increased at 50 and 250 μM Cd, unchanged at 1 mM Cd, and decreased at 5 mM Cd (Geiken et al., 1998). At 20 μM and 80 μM Cd, NPQ decreased due to the reduction of fast relaxing component qE; simultaneously, its slow relaxing component qI increased (Lysenko et al., 2015). In these studies, non-photochemical quenching was presented as a single-point measurement after a light induction. The data in Fig. 3A,B shows nonlinear dynamics; we can describe different Cd effects selecting particular points from these dynamics. It is favorable showing results as a dynamic process obtained with the use of different AL; Cd effect should be studied as a combination of IC and RLC or several IC at different AL intensities.

#### 4.3.2 Photochemical quenching

The decrease of non-photochemical quenching retained more light energy in PSII. This extra energy was distributed between open (X(II)) and closed (qC) PSII. The pattern of changes was diverse. In IC measured at AL 128 μmol photons m^-2^ s^-1^, the changes of non-photochemical quenching (Fig. 3A,B) mostly influenced open PSII both up and down (Fig. 2A,B, Fig. 4A,B). In RLC (Fig. 3C,D), the decrease of qN mostly resulted in the increase of closed PSII while the increase of qN mainly resulted in the decrease of open PSII (Fig. 2C,D, Fig. 4C,D).

The balance between open and all (open+closed) PSII (qP) was nearly unchanged in maize at 80 μM Cd; in other cases, qP decreased (Fig. 2E-H). In barley under both Cd concentrations, qP was decreased due to qC increase. During first 40-120 s since the light induction, Cd treatment increased both X(II) (Fig. 2A) and qC (Fig. 4A) whereas qP remained unchanged (Fig. 2E); no qP changing indicates that both increases were proportional to their ratio in the control. Since 160 s, X(II) was increasing in the control and reached the level of Cd-treated variants (Fig. 2A), while the small difference of qC between the control and Cd-treated variants remained relatively stable (Fig. 4A). The decreasing qP in Cd-treated plants (Fig. 2E) reflects losing of difference in X(II) and persisting difference of qC. In RLC, the impact of qC was obvious (Fig. 2C,G, Fig. 4C). In maize at 250 μM Cd, qP was decreased due to X(II) decrease. Thus, Cd treatment induced a complicated pattern of changes. The alterations of photochemically active PSII cannot be explained within the framework of this experimental model.

In barley plants, Cd stress accelerated readaptation of the photochemical activity of PSII to the light conditions. Cd-treated plants reached a stationary level faster than in the control (Fig. 2A). In maize plants at 80 μM Cd, the similar acceleration was also observed; however, it was very small (Fig. 2B). The similar accelerative effect was observed for PSI (see section 4.2). In barley plants, heat stress accelerated the restoration of X(II) during the first two minutes after AL induction (Lysenko et al., 2023). The accelerated readaptation of the photosynthetic electron transport chain to light conditions can be a general feature of diverse abiotic stresses that was not observed before. This accelerative effect was more specific to barley young plants than to maize ones.

#### 4.3.3 Acceptor-side limitation of PSII (qC)

The observed changes in closed PSII can be explained by a two-component mechanism. At 80 μM Cd, qC changed in agreement with Cd accumulation in the chloroplasts. Barley chloroplasts accumulated a large amount of Cd while maize chloroplasts accumulated small portion of Cd (Lysenko et al., 2015). In RLC under rather high AL intensities, qC demonstrated a large increase in barley plants and a small increase in maize plants (Fig. 4C,D). In IC under relatively low AL intensity, qC increased slightly in barley and remained practically unchanged in maize (Fig. 4A,B). Thus, we can suppose a direct Cd effect on qC under 80 μM Cd.

Increasing the concentration from 80 μM to 250 μM Cd reversed this effect. The level of qC decreased at 250 μM comparing with 80 μM Cd (Fig. 4); the decrease was similar in barley and maize. The differences between the mean qC values at 80 μM and 250 μM Cd are shown in Fig. 10. The concentration shift from 80 μM to 250 μM Cd induced diverse changes of Cd content in the chloroplasts and a similar increase of general toxicity in barley and maize plants (Lysenko et al., 2015). Thus, the similar decreases (Fig. 10) should be referred to an indirect effect of Cd at 250 μM. We hypothesize that the portion of closed PSII was increased by means of a direct mechanism at 80 μM Cd; at 250 μM Cd, this effect was overcome via an indirect mechanism decreasing qC.

**Fig. 10.**
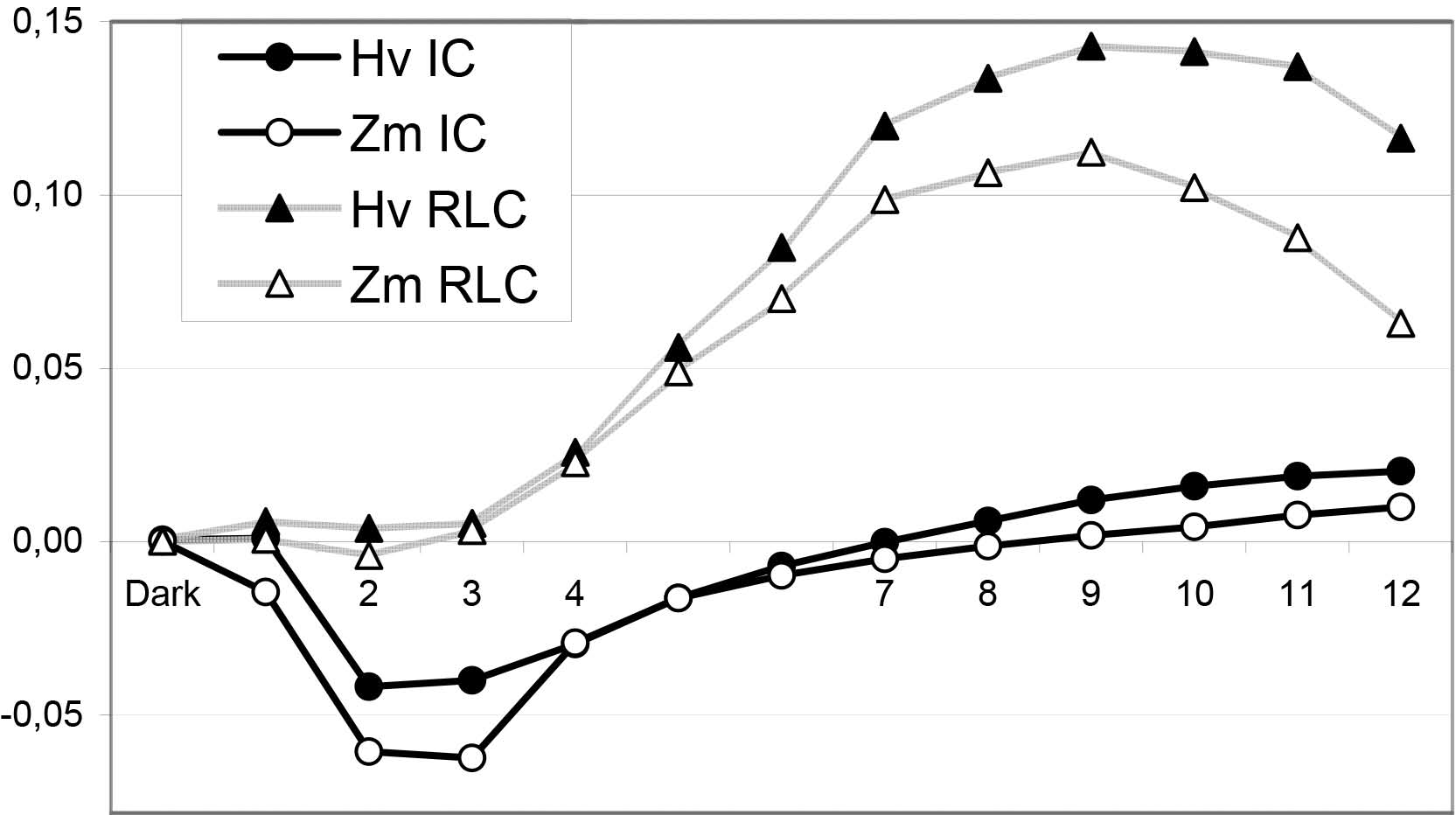
The impact (Δ) on qC of increasing Cd concentration from 80 μM to 250 μM (from Fig. 4). The means from Fig. 4 were recalculated to show effect of the concentration increase: ΔqC_t_ = qC_t_ (Cd80) - qC_t_ (Cd250). Positive values reflect the decreasing effect of 250 μM; negative values denote the increasing effect. Black lines with circles indicate IC; grey lines with triangles represent RLC. Filled symbols show barley (Hv); open symbols show maize (Zm). The abscissa numbering indicates both time and AL intensities. For IC, 1-12 correspond to the range 0.5 - 440 s; for RLC, 1-12 correspond to the range 36 – 1954 μmol photons m^-2^ s^-1^. The digits 1, 5, and 6 are omitted for better visibility.

#### 4.3.4 Possible mechanisms

The direct inhibitory action of Cd on the oxygen-evolving complex of PSII is generally accepted. It was shown both *in vitro* (Bazzaz and Govindjee, 1974; Sigfridsson et al., 2004) and *in vivo* (Baszynski et al., 1980). In barley chloroplasts, the quantity of Cd was sufficient to replace one of the four Mn in the oxygen-evolving complex (Lysenko et al., 2019). Artificial water-oxidizing complexes consisted of one, two, or four Mn atoms; some artificial Mn-catalysts comprised Cd^2+^ as the redox-inert ion stabilizing a crystal structure (reviewed in Najafpour et al., 2016). Therefore, we are unsure whether a single Cd/Mn substitution is capable to inhibit Mn_4_CaO_5_ cluster or not. Anyway, this mechanism cannot explain the changes of PSII activity observed in the current study (Fig. 1-4).

Otherwise, the hypothesis of Cd/Cu substitution in plastocyanin (see section 4.2) explains the inhibition of electron transfer between the acceptor-side of PSII and donor-side of PSI. The bottleneck in electron traffic can increase the portion of PSII reaction centers limited (closed) at the acceptor-side at 80 μM Cd that was masked with an indirect mechanism at 250 μM Cd.

Phosphate shortage can limit the photosynthesis by restricting ATP synthesis (Sage and Kubien, 2007). Suppressing ATP-synthase activity inhibits unloading of ΔpH gradient and maintaines a higher level of pH-dependent non-photochemical quenching. In the late part of IC, increasing Cd concentration slowed down the decay of non-photochemical quenching. In both species, the angles of qN and NPQ decay slopes ranged similarly: control > 80 μM Cd > 250 μM Cd (Fig. 3A,B, Suppl. Fig. S3A,B). Consequently, an indirect Cd action inhibited exhausting of ΔpH gradient, probably, by limiting the ATP-synthase activity. Cd impact on inorganic phosphate in chloroplasts had not been studied. However, the phosphate shortage may be the indirect mechanism of Cd action that decreased qC and overrode the direct effect under 250 μM Cd (Fig. 10). We suppose the next chain of consequences: phosphate shortage → ATP-synthase inhibition → slowdown of ΔpH exhaustion → increasing of non-photochemical quenching → decreasing of light energy in PSII and portion of closed PSII (qC).

### 4.4. Balance between activities of PSI and PSII

We compared the photochemical activities of PSI (Y(I)) and PSII (X(II)) under illumination. After the light induction, the ratio Y(I)/X(II) gradually increased in control variants of both species; the ratio reached the maximal level of about 2.5 in 2 min and further decreased slowly to the level of about 2 (Fig. 7A,B). Cd treatment accelerated the readaptation to light conditions in barley plants. The ratio Y(I)/X(II) reached the steady state level very fast (0.5 s), though this level was lower than in the control (Fig. 7A). This acceleration was specific to barley and Cd treatment (Fig. 7A,B). Thus, Cd-treated barley plants readapted the photochemical activities of PSI, PSII, and their balance fast after the light induction. Under heat stress, Y(I)/X(II) also increased faster during the first second since the light induction (Lysenko et al., 2023). However, this increase was smaller and observed in both barley and maize plants; as a whole, the increases of Y(I)/X(II) induced with Cd and heat stresses were dissimilar.

Cd treatment decreased the ratio Y(I)/X(II) in barley plants in both IC and RLC (Fig. 7A,C). This means larger inhibition of PSI comparing with PSII that was concluded previously (Lysenko et al., 2020). The small decrease of Y(I)/X(II) was observed in maize at 80 μM Cd; the decrease was significant at some points of IC and RLC (Fig. 7B,D). The small size of the decrease can be attributed to the small Cd content in maize chloroplasts at 80 μM Cd (Lysenko et al., 2015). However, Y(I)/X(II) was increased in maize plants at 250 μM Cd; it was observed in both IC and RLC (Fig. 7B,D). In this case, we should conclude that PSII was inhibited more than PSI.

The inhibitory effect of 250 μM Cd was calculated for both Y(I) and X(II) from the mean values (Fig. 11). The size of decrease was similar in Y(I) and X(II); in many points, Y(I) was reduced more than X(II). However, the values of Y(I) (Fig. 5B,D) were much larger than the values of X(II) (Fig. 2B,D); therefore, Y(I) were reduced proportionally smaller comparing with X(II) (Fig. 7B,D). For leaves with the single type mesophyll chloroplasts, we have to follow proportional changes of Y(I) and X(II) that, probably, reflects some stoichiometry and/or relative activities of PSI and PSII. However, maize is C_4_ plant with typical Kranz anatomy; these plants bear nearly equal amounts of mesophyll and bundle sheath cells in their leaves (Langdale, 2011). Mesophyll cells include chloroplasts with both PSII and PSI; in C_4_ plants, bundle sheath cells contain chloroplasts with PSI and no or very small activity of PSII (Turkan et al., 2018). In maize, PSII can be inhibited practically in mesophyll type chloroplasts only. Whether PSI was inhibited in one or both types of chloroplasts?

**Fig. 11.**
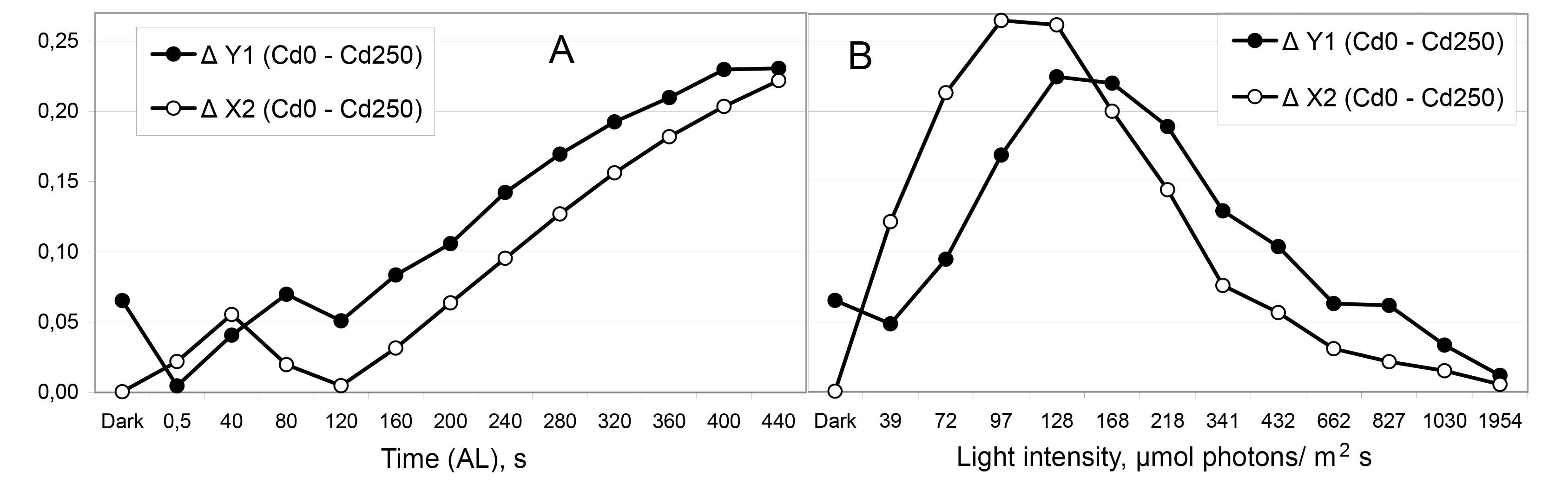
The impact (Δ) of 250 μM Cd on the photochemical activities Y(I) of PSI and X(II) of PSII in maize plants (from Figs. 2, 5). Corresponding means from Fig. 2B,D and Fig. 5B,D were recalculated to show the effect of 250 μM Cd comparing with the control (0 μM Cd): ΔY(I)_t_ = Y(I)_t_ (Control) - Y(I)_t_ (Cd250); ΔX(II)_t_ = X(II)_t_ (Control) - X(II)_t_ (Cd250). Positive values reflect the inhibitory effect of 250 μM Cd. A – IC; B – RLC. Filled symbols show ΔY(I); open symbols show ΔX(II).

First, we do not know Cd content in bundle sheath cell chloroplasts. Our previous study was focused on mesophyll chloroplasts; therefore, the leaf homogenization was rather mild (Lysenko et al., 2015). The destruction of bundle sheath cells for chloroplast isolation requires a harder procedure (Jenkins and Boag, 1985; Romanowska and Parys, 2011). Therefore, we have to consider three possibilities. A) In both types of chloroplasts, PSI activity decreased similarly; so, PSI was inhibited less than PSII in the mesophyll chloroplasts. B) In the mesophyll chloroplasts, PSI and PSII were inhibited similarly while PSI remained unchanged in the bundle sheath cell chloroplasts. C) In the mesophyll chloroplasts, PSI was inhibited more than PSII; however, PSI activation in the bundle sheath cell chloroplasts compensated it. The question of Cd action on processes in bundle sheath cell chloroplasts has not been previously analyzed. Therefore, we cannot decide which of the variants is true.

Second, the similar effect was observed under heat stress. Under the most severe conditions (46°C at low air humidity), X(II) decreased more than Y(I) that increased the ratio Y(I)/X(II) in maize plants (Lysenko et al., 2023). Barley plants demonstrated no increase of Y(I)/X(II) and died just after completing the experiment while maize plants survived (Lysenko et al., 2023). These data interfere with considering large X(II) decrease under severe stress as an inhibitory process. Can we find an adaptive explanation for decreasing PSII activity?

Plants with C_4_ type of photosynthesis (including maize) have a lesser Rubisco content (Ku et al., 1979) and smaller proportion of Rubisco oxygenase activity (Turkan et al., 2018) than C_3_ plants. Elevating temperature enhances the impact of oxygenase activity by reducing Rubisco affinity to CO_2_ and increasing its affinity to O_2_ (Laing et al., 1974). Cd treatment decreased Rubisco content (Pietrini et al., 2003; Lysenko et al., 2015). As a C_4_ plant, maize features restricting PSII-dependent oxygen production and enhancing energy acquisition from the cyclic electron transport around PSI (e.g. Fig. S1A). Considering this, the adaptive explanation can be suggested as follows.

Under very severe stress (250 μM Cd, 46°C), maize plants upregulated non-photochemical quenching (Fig. 3B,D) to restrict PSII activity (Fig. 2B,D), PSII-dependent oxygen production and oxygenase activity of Rubisco. The possible upregulating mechanism was discussed in section 4.3.4. Restricting PSII-dependent O_2_ production can increase ribulose-1,5-bisphosphate carboxylation and reduce both the production of toxic 2-phosphoglycolate and waste of energy for its inactivation through the mechanism of photorespiration (Langdale, 2011; Turkan et al., 2018). Under conditions of heavy stress, declining Rubisco oxygenase activity can be more advantageous comparing with energy acquisition from PSII.

### 4.5. Balance between limitations of PSI and PSII

The ratios of the limitations were calculated for the first time; so, we will consider some details. The ratio qC/Y(NA) shows the balance of limitations at the acceptor sides of PSII (qC) and PS(I) (Y(NA)). Half a second since the light induction, both limitations were large (Fig. 4A,B, Fig. 6E,F). In maize, both limitations were nearly equal (Fig. 8B inset); in barley, qC was 15% smaller than Y(NA) (Fig. 8A inset). Cd treatment had no effect in maize and equalized both limitations in barley; the latter effect was specific to Cd (Fig. 8A,B insets). In 2 min, qC/Y(NA) reached a peak followed by a decrease (Fig. 8A,B). In barley, Cd treatment accelerated reaching the peak, which was not specific to Cd (not shown). The steady state level was reached in 4 min in barley. In maize, control plants demonstrated no peak; in Cd-treated plants, slow decreasing ratio reached the control level at the end of measurement (7.5 min).

In the control, both limitations were approximately equal or qC was a little larger than Y(NA) (qC/Y(NA) ≈ 1.0-1.3-1.5); in rare cases, qC was substantially smaller or larger than Y(NA) (Fig. 8). Under Cd treatment, qC was mostly larger than Y(NA); the difference had reached 5 times. At the steady state level, qC was two times larger than Y(NA) in barley (Fig. 8A). Once, qC was substantially smaller than Y(NA) (Fig. 8C). Therefore, the photochemical activity of PSII was not limited by Calvin-Benson cycle. In the control, limitations of the electron-transport chain and Calvin-Benson cycle were balanced; in Cd-treated plants, limitation of the electron-transport chain prevailed. In most parts, the dynamics of qC/Y(NA) (Fig. 8) simply reflects the changes of qC (Fig. 4).

The ratio qC/Y(ND) enables estimating limitations between the acceptor side of PSII (qC) and the donor-side of PS(I) (Y(ND)). It is a tool for studying the fragment of electron-transfer chain including plastoquinone pool, cytochrome b_6_/f, and plastocyanin.

On the contrary, qC was larger than Y(ND) at the beginning of the curves solely (Suppl. Fig. S6). In IC, qC > Y(ND) in maize plants and control barley plants after 40 s since the light induction; qC ≈ Y(ND) in Cd-treated barley plants after 40 s and in control maize plants after 80 s yet (Suppl. Fig. S6A,B, Fig. 9). In the beginning of RLC, control plants demonstrated qC > Y(ND) in barley and qC ≈ Y(ND) in maize (Suppl. Fig. S6C,D). In the most parts of the curves, qC was smaller than Y(ND). Consequently, this segment of the electron-transport chain was limited mostly at the donor side of PSI.

Cd treatment increased the ratio qC/Y(ND) in barley, mostly unchanged it in maize at 80 μM Cd, and decreased it in maize at 250 μM Cd (Suppl. Fig. S6; Fig. 9). The changes of both ratios qC/Y(NA) and qC/Y(ND) cannot be explained within the framework of this experiment.

Surprisingly, we found that short quasi-stationary levels were characteristic feature of qC/Y(ND) dynamics (Fig. 9). The values of quasi-stationary levels varied from 0.2 to 0.65 (qC < Y(ND)). In most cases, one variant showed equal values of quasi-stationary levels in both IC and RLC; in barley at 80 μM Cd, they differed substantially. In a variant, IC and RLC reached equal quasi-stationary levels in similar (Fig. 9B) or different time (Fig. 9A). The quasi-stationary levels did not reflect mutual plateaus; instead, they were found in actively changing segments of qC and Y(ND) dynamics. Three examples are shown in Fig. 12. In IC, the quasi-stationary levels were followed by the increases due to further decreases of Y(ND) (Fig. 6A,B). In RLC, the quasi-stationary levels were followed by the decreases mostly due to the decreases of qC (Fig. 4C,D) because of the corresponding increases of non-photochemical quenching (Fig. 3C,D).

**Fig. 12.**
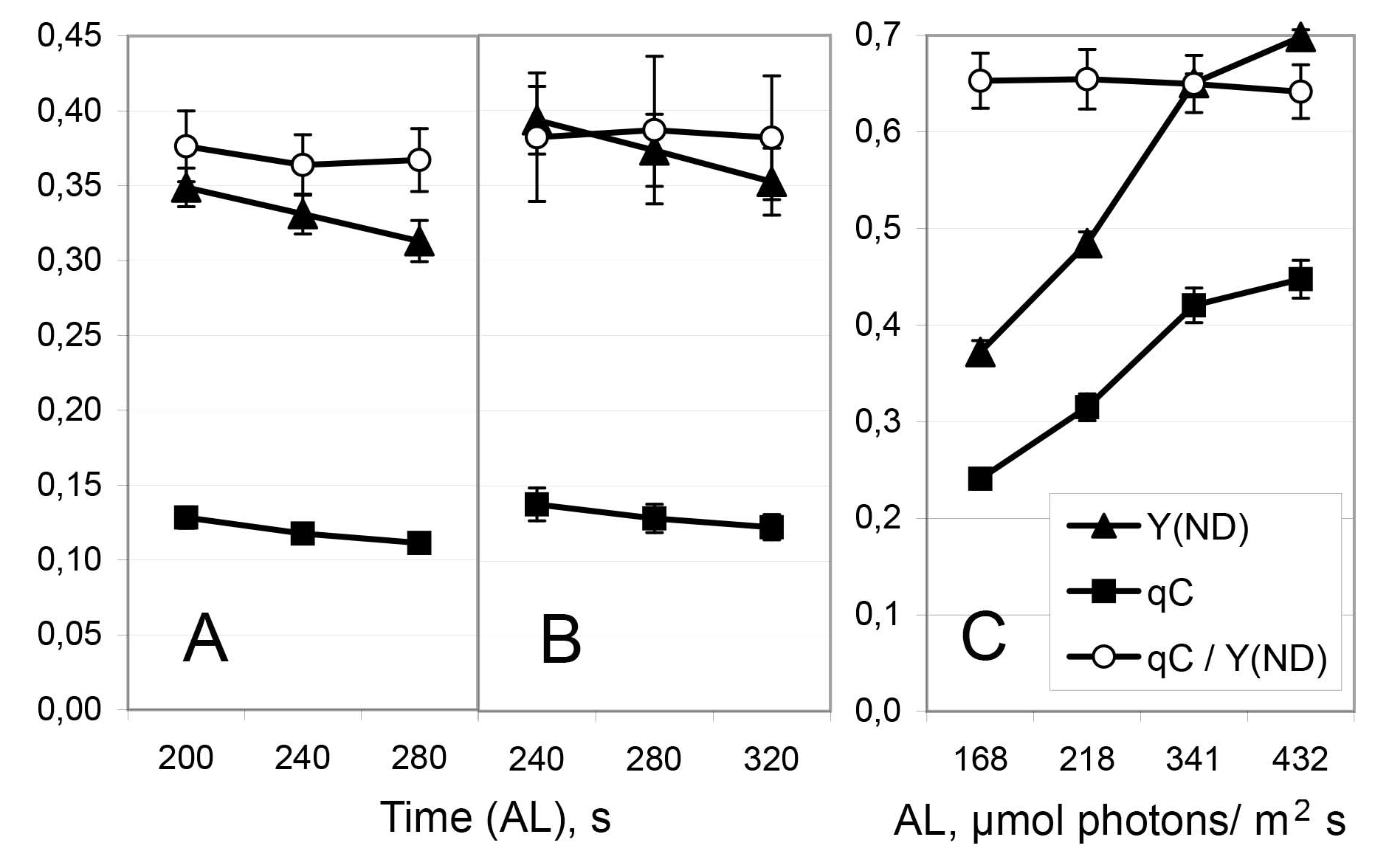
The segments of qC, Y(ND), and qC/Y(ND) dynamics are shown in one figure. Selected examples demonstrating quasi-stationary levels of the ratio qC/Y(ND). A – IC control maize (from Figs. 4B, 6B, 9B/S6B); B - IC Cd80 maize (from Figs. 4B, 6B, 9B/S6B); C – RLC Cd80 barley (from Figs. 4C, 6C, 9C/S6C). Filled squares – qC; filled triangles – Y(ND); open circles – the ratio qC/Y(ND). Means ± SE.

We hypothesize that the quasi-stationary levels reflected simultaneous proportional changes of qC and Y(ND) as the result of an unknown regulatory process. In stationary segments, Y(ND) > qC. Therefore, we consider limitation at PSI donor-side as the primary event. Limitation at the donor side of PSI should increase redox state of plastocyanin, cytochrome b_6_/f, and plastoquinone pool. Sensing of plastoquinone redox state is the basis of regulatory processes in chloroplasts (Tullberg et al., 2000; Mattila et al., 2020). Probably, barley and maize chloroplasts recognize a bottleneck in electron traffic to PSI through oversaturation of plastoquinone pool and induce unloading of closed PSII by increasing non-photochemical quenching. Comparing to PSI donor-side limitation, limitation of PSII at the acceptor side can be decreased by one third (Fig. 9C) or by five times (Fig. 9A). The depth of this fall depends on particular plant species under particular conditions. We realize the importance of this finding. The simultaneous proportional changes of qC and Y(ND) have to be thoroughly analyzed in a separate manuscript.

## 5. Conclusion

In young barley and maize plants, Cd treatment altered the activities of PSII and PSI in different way. Both maximal and relative photochemical activities of PSI changed in agreement with Cd accumulation in the chloroplasts of these plants. The photochemical activity of PSI was mostly limited at the donor side; the changes of PSI donor-side limitation also mirrored Cd accumulation in the chloroplasts. The observed inhibition of PSI can be explained by the direct Cd substitution of Cu in plastocyanin. The observed changes in the photochemical and non-photochemical quenching by PSII can be clearly explained by neither a direct nor an indirect Cd action. Acceptor-side limitation of PSII can be explained with a complex mechanism. At 80 μM Cd, PSII was limited at the acceptor side through a direct Cd action; at 250 μM Cd, this effect was overcome by an indirect Cd action and closed PSII were unloaded, probably, by means of non-photochemical quenching.

We compared the limitations at the acceptor-side of PSII (qC), donor-side of PSI (Y(ND)), and acceptor-side of PSI (Y(NA)). In general, these limitations were ranged as follows: Y(NA) < qC < Y(ND). Short sections of IC and RLC dynamics demonstrated proportional changes of qC and Y(ND). This indicates existence of unknown mechanism adjusting qC value to a size of Y(ND). Therefore, acceptor-side limitation of PSII can also be explained by Cd/Cu substitution in plastocyanin; it was obvious at 80 μM Cd and masked by an indirect mechanism at 250 μM Cd. A phosphate shortage in chloroplasts may be hypothesized as an indirect mechanism decreasing qC.

## Author contribution

**Eugene Lysenko:** Conceptualization, Methodology, Investigation, Visualization, Formal Analysis, Writing; **Victor Kusnetsov:** Funding Acquisition, Resources. All authors read and approved the manuscript.

## Supporting information

Supplementary Tabs and Figs

## Abbreviations

AL: actinic light
Chl: chlorophyll
IC: induction curve
PAM: pulse amplitude modulation
PSI, PSII: photosystem I and II
P_700_: reaction center Chl of PSI
RLC: rapid light curve
SE: standard error
SP: saturation pulse Methods 1+6 (1 page)

## PAM terms

Fv/Fm: maximal quantum yield of PSII
Փ_PSII_: effective quantum yield of PSII
qP: photochemical coefficient of Chl fluorescence
qN, NPQ: coefficients of non-photochemical quenching of excited Chl state
qC: coefficient of acceptor-side limitation of PSII (closed PSII)
X(II): relative quantum yield of PSII
Y(ND): coefficient of donor-side limitation of PSI
Y(NA): coefficient of acceptor-side limitation of PSI
Fo / Fo': minimal Chl fluorescence in dark / light
Fm / Fm': maximal Chl fluorescence in dark / light
Fv / Fv': variable Chl fluorescence in dark / light
Fs: steady state Chl fluorescence in light
Po: minimal light absorption level
Pm / Pm': maximal P_700_ change in dark / light
P: steady state P_700_ absorption level in light

## Acknowledgement

The authors thank Dr. A.A. Klaus for the assistance in plant growing. The seeds of maize were kindly provided by Dr. E.V. Kartamysheva (VNIIMK, Don experimental station). The research was carried out within the state assignment of Ministry of Science and Higher Education of the Russian Federation (theme No. 122042700044-6) and supported by the Russian Science Foundation grant (project No. 14-14-00584).

