## Supplementary Tabs and Figs for "Changes of Cd content in chloroplasts are mirrored by the activity of photosystem I, but not by photosystem II"

**Source:** bioRxiv

### General map of deleted points for calculating the ratios Y(I)/X(II), qC/Y(NA), and qC/Y(ND).

Table S1. The ratio Y(I)/X(II). Map of points deleted because of small denominator value.

| | Barley | | | Maize | | |
|---|---|---|---|---|---|---|
| | Control | Cd 80 | Cd 250 | Control | Cd 80 | Cd 250 |
| Number of IC: | 17 | 17 | 37 | 18 | 18 | 24 |
| Dark | | | | | | |
| 0,5 | | | 3 | | | 1 + 1 |
| 40 | | | | | | |
| 80 | | | | | | |
| 120 | | | | | | |
| 160 | | | | | | |
| 200 | | | | | | |
| 240 | | | | | | |
| 280 | | | | | | |
| 320 | | | | | | |
| 360 | | | | | | |
| 400 | | | | | | |
| 440 | | | | | | |
| Number of RLC: | 17 | 17 | 21 | 18 | 18 | 19 |
| 39 | | | | | | |
| 72 | | | | | | |
| 97 | | | | | | |
| 128 | | | | | | |
| 168 | | | | | | |
| 218 | | | | | | |
| 341 | | | | | | |
| 432 | | | | | | |
| 662 | 1 * | | | | | |
| 827 | 1 * | | | | | |
| 1030 | | | | | | |
| 1954 | | 1 | | | | 1 |

Deleted X(II): negative value; too small values (maize 0.001; barley 0.004-5).

* - data absent.

Table S2. The ratio qC/Y(ND). Map of points deleted because of small denominator value.

| | Barley | | | Maize | | |
|---|---|---|---|---|---|---|
| | Control | Cd 80 | Cd 250 | Control | Cd 80 | Cd 250 |
| Number of IC: | 17 | 17 | 37 | 18 | 18 | 24 |
| 40 | 1 + **1** + 2 | | | | | |
| 80 | | | | | | |
| 120 | | | | | | |
| 160 | | | | | | |
| 200 | | | | | | |
| 240 | | | | | | |
| 280 | | | | | | |
| 320 | | | | | | |
| 360 | | | | | | |
| 400 | | | | | | |
| 440 | | | | | | |
| Number of RLC: | 17 | 17 | 21 | 18 | 18 | 19 |
| 72 | 2 + **7** | **2** | 1 + **2** | 1 + **1** | **1** + 1 | |
| 97 | **4** | | | **1** | | |
| 128 | | | | | | |
| 168 | | | | | | |
| 218 | | | | | | |
| 341 | | | | | | |
| 432 | | | | | | |
| 662 | 1 * | | | | | |
| 827 | 1 * | | | | | |
| 1030 | | | | | | |
| 1954 | | | | | | |

Deleted Y(ND): negative value; zero value (0); **value ≤ 0.025**; value ≥ 0.030.
* - data absent.

Table S3. The ratio qC/Y(NA). Map of points deleted because of small denominator value.

| | Barley | | | Maize | | |
|---|---|---|---|---|---|---|
| | Control | Cd 80 | Cd 250 | Control | Cd 80 | Cd 250 |
| Whole IC deleted: | 1 | 2 | 1 | | | |
| Number of IC: | 16 | 15 | 36 | 18 | 18 | 24 |
| 0,5 | | | | | | |
| 40 | | | **1** | | | |
| 80 | **1** | | **2** | | **1** | **1** |
| 120 | | | 1 + **3** | | | 2 |
| 160 | | | 3 + **2** | | **1** | 1 + 1 |
| 200 | **1** | | 2 + **1** | | | |
| 240 | | **1** | 2 + **1** | | | |
| 280 | **2** | | 1 + **2** | | | 1 + **2** |
| 320 | | **1** + **1** | **2** | | | |
| 360 | | | **2** | | | |
| 400 | | | **2** | | | **2** |
| 440 | | **1** | 1 + **2** | | **2** | 1 + **2** |
| Whole RLC deleted: | 1 | 1 | 1 | | | |
| Number of RLC: | 16 | 16 | 20 | 18 | 18 | 19 |
| 72 | | **1** | 1 + **1**+ 1 | | | 1 |
| 97 | | | | | | **1** |
| 128 | | | **1** | | | 1 |
| 168 | **1** | | | | **1** | **1** |
| 218 | **1** | | 1 | | | |
| 341 | | | | | | **2** |
| 432 | | | 1 | | | |
| 662 | | | | | | |
| 827 | | **1** | 1 | | | |
| 1030 | | 1 | | | | |
| 1954 | | | | | **1** | |

Deleted Y(ND): negative value; zero value (0);
**value ≤ 0.025**; **0.025 < value > 0.030**; value ≥ 0.030.

Fig. S1. Y(I) and X(II) in darkness and 0.5 s after the light induction.

The corresponding values from Fig. 2A,B and Fig. 5A,B are shown with the higher resolution.

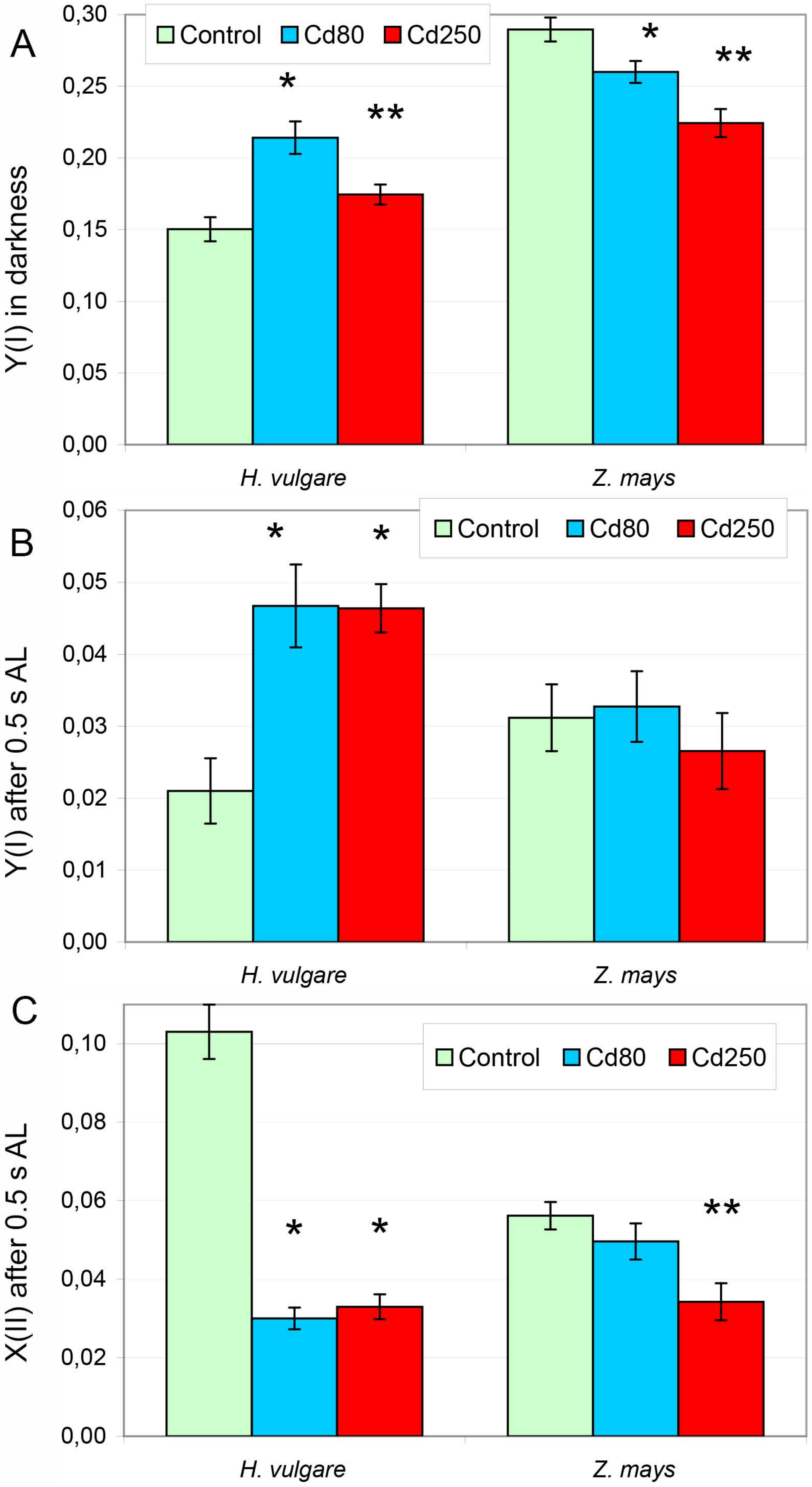


Means ± SE.

* - the difference from control is significant ($p \leq 0.05$).

** - the difference from both control and Cd 80 µM is significant ($p \leq 0.05$).

Fig. S2. Dynamics of photochemical coefficient $\Phi_{PSII}$ of PSII.

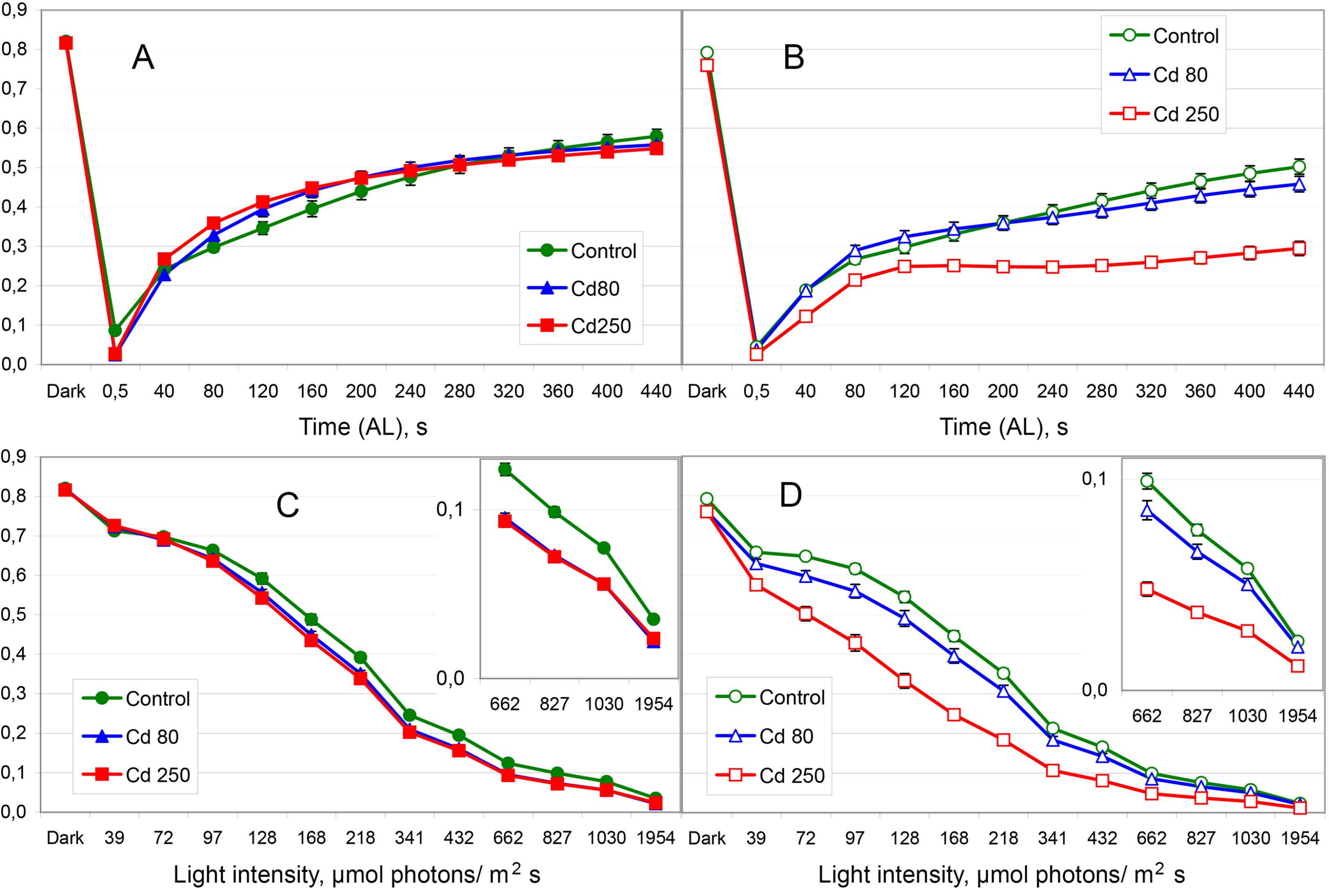


A, B – IC; C, D – RLC. A, C – barley; B, D – maize. Means ± SE. All designations are the same as in Fig. 2.

Fig. S3. Dynamics of non-photochemical quenching of PSII: coefficient NPQ.

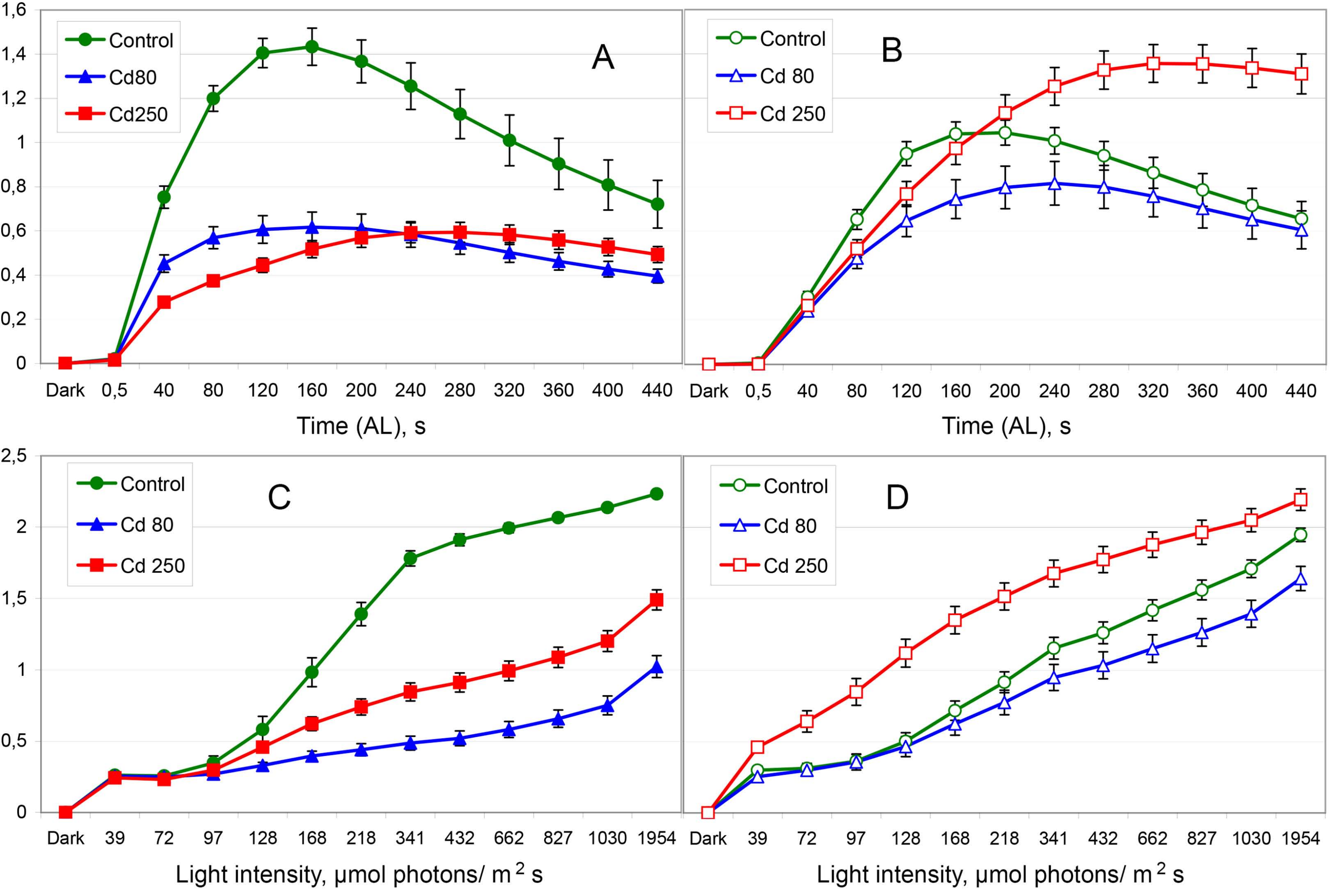


A, B – IC; C, D – RLC. A, C – barley; B, D – maize. Means ± SE. All designations are the same as in Fig. 2.

### Fig. S4. Dynamics of qN.

Data from Fig. 3 panels C and D are presented together for the comparison. (0 = Control)

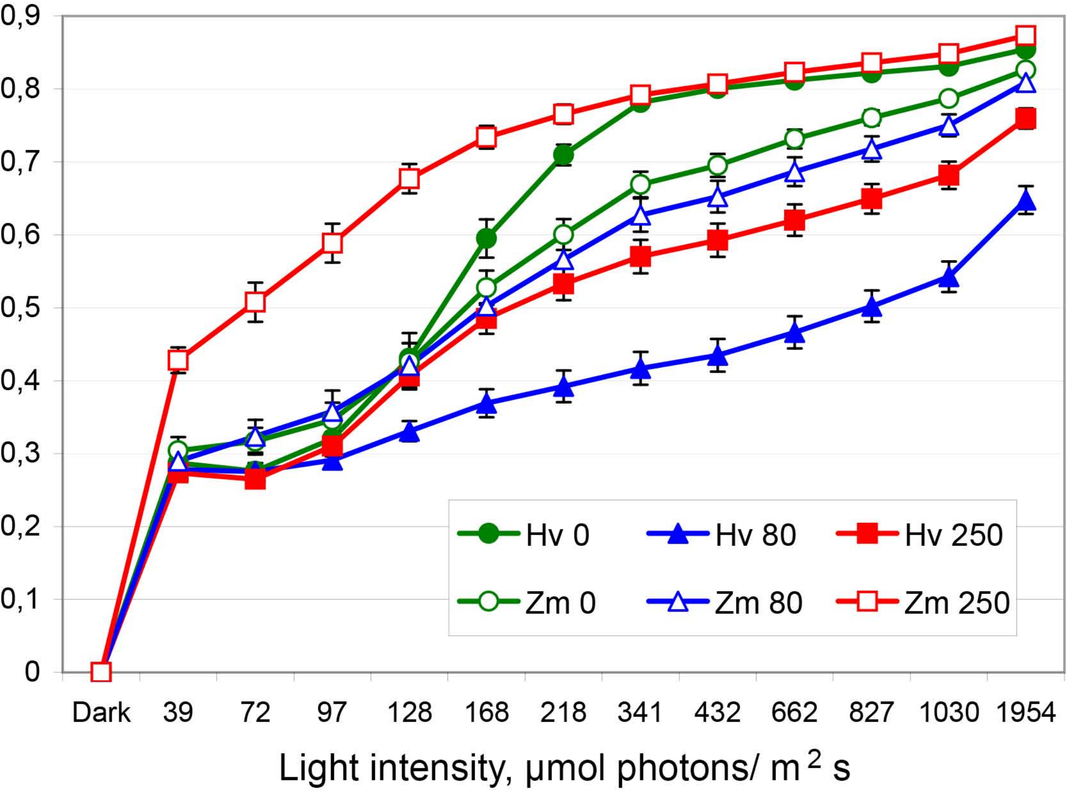


### Fig. S5. Dynamics of Y(I)/X(II).

Data from Fig. 7 panels C and D are presented together for the comparison. (0 = Control)

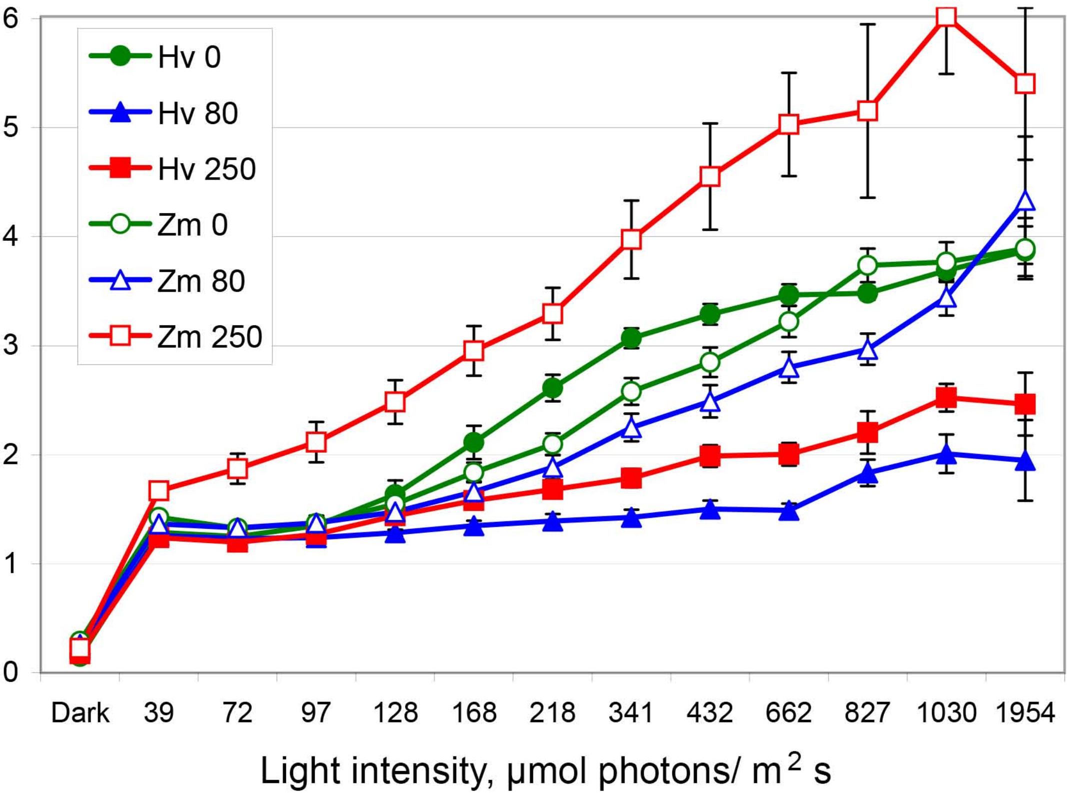

Fig. S6. The ratio qC/Y(ND) showing balance of limitations between the acceptor side of PSII (qC) and the donor side of PSI (Y(ND)). Complete data set.

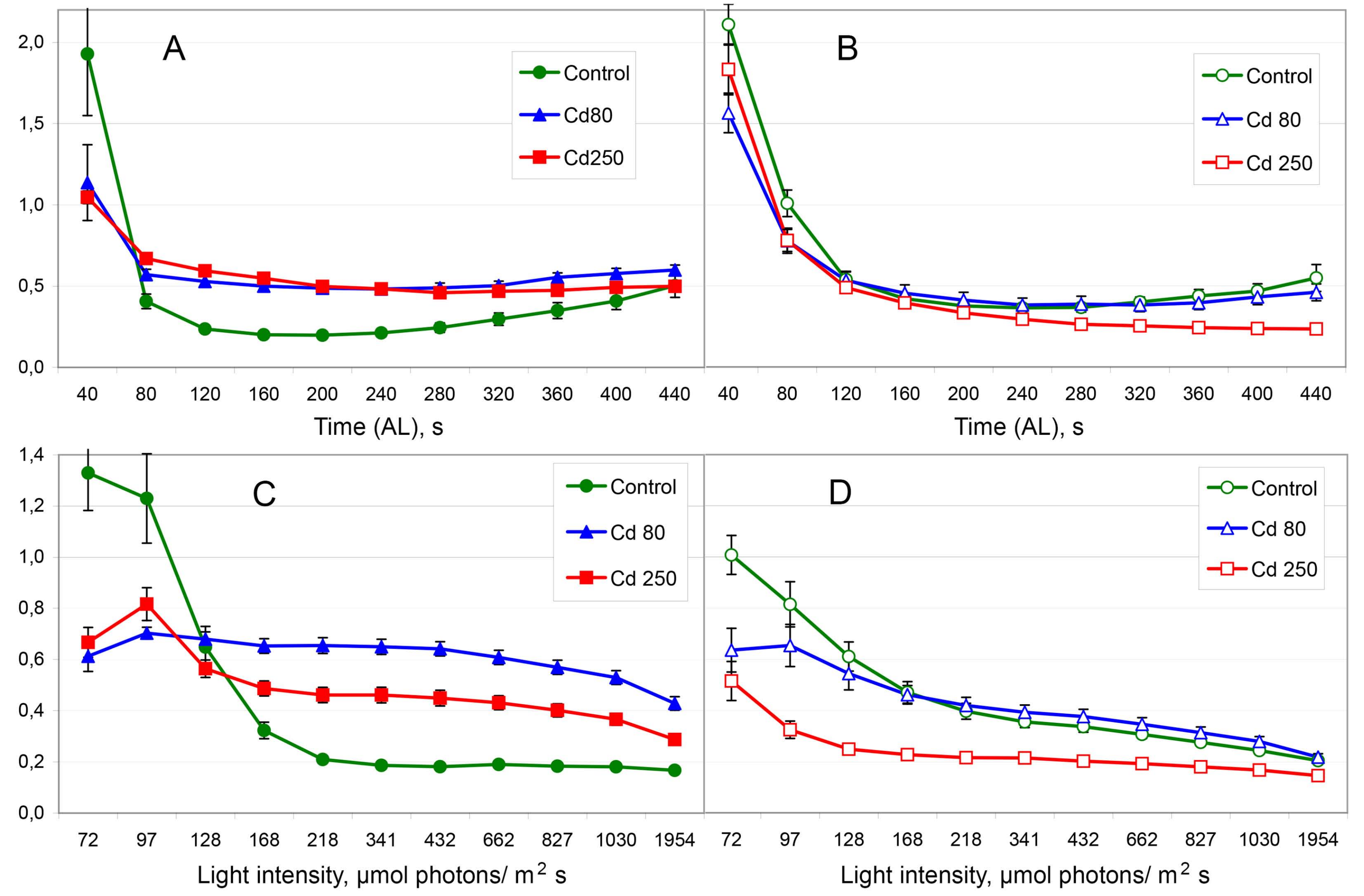


A, B – IC; C, D – RLC. A, C – barley; B, D – maize. Means ± SE. All designations are the same as in Fig. 2.

Fig. S7. IC dynamics of Fo.

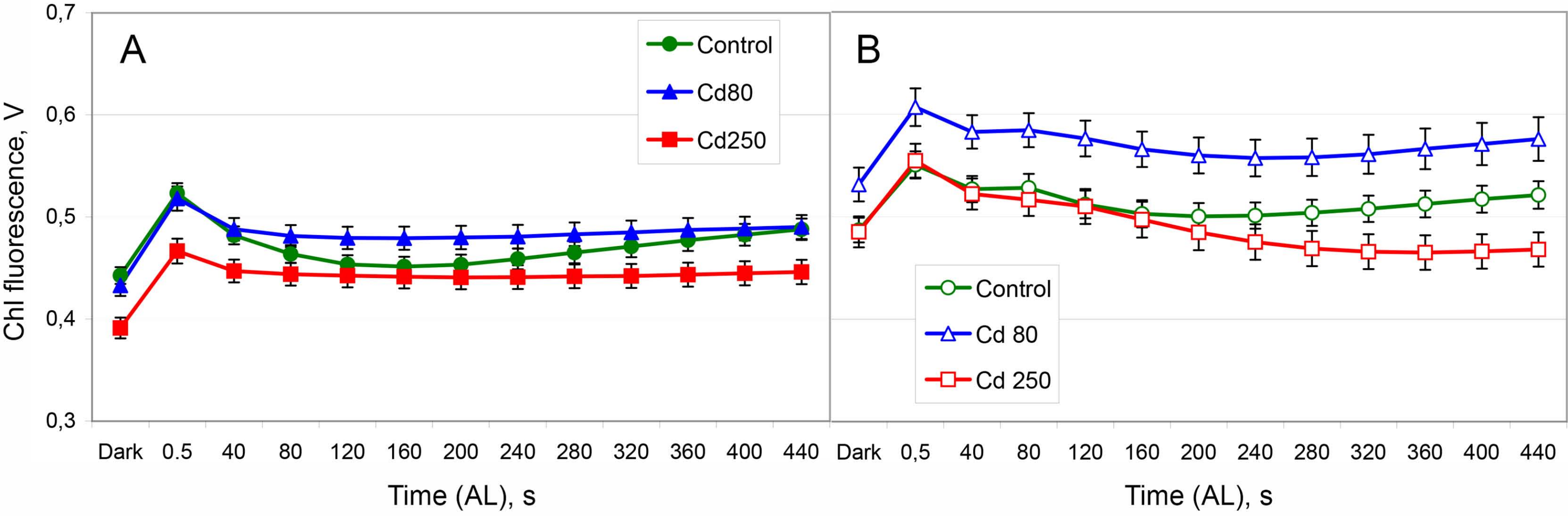


A – barley; B – maize. Means ± SE. All designations are the same as in Fig. 2
